# NPAS4 refines spatial and temporal firing in CA1 pyramidal neurons

**DOI:** 10.64898/2026.03.22.713468

**Authors:** Anja Payne, Daniel A. Heinz, Chiaki Santiago, Lara L. Hagopian, Rolando Sceptre Ganasi, Clare Quirk, Andrea L. Hartzell, Jill K. Leutgeb, Stefan Leutgeb, Brenda L. Bloodgood

## Abstract

NPAS4 is an activity-dependent transcription factor that, in CA1 of the hippocampus, regulates inhibitory synapses made onto the active pyramidal neuron. In principle, NPAS4 thereby allows the past activity of a neuron to influence how it encodes information, although this has not yet been demonstrated. Here, we generated a sparse, CA1-specific knockout (KO) of NPAS4 in the mouse hippocampus and used optogenetic tagging to identify KO neurons *in vivo*. Recordings from intermingled wild-type (WT) and KO neurons in awake behaving animals revealed that NPAS4 deletion degrades spatial representations and temporal precision of spiking: KO neurons exhibited larger place fields with reduced in-field firing and increased out-of-field firing, less stable place fields, reduced coupling to local field potential theta oscillations, and diminished phase precession. These findings demonstrate that NPAS4 plays a crucial role in refining the spatial and temporal properties of CA1 pyramidal neuron spikes, which themselves are thought to be fundamental building blocks of more complex processes such as learning and memory.

## INTRODUCTION

Experiences drive long-lasting changes in brain function through a range of molecular mechanisms, including the induction of activity-dependent transcription factors. These transcription factors are rapidly induced by transient neuronal activity and initiate gene expression programs that result in persistent alterations to neuronal function and synaptic connectivity^1–6^. Although a growing body of research has linked the expression of activity-dependent transcription factors to learning, memory, and behavior^7–12^, relatively few studies have directly examined how these factors influence the encoding of information in awake, behaving animals^13–17^.

In mice, the transcription factor NPAS4 is expressed almost exclusively in neurons and can be induced across multiple brain regions in a stimulus-dependent manner^13,18–25^. For instance, contextual fear conditioning or exposure to an enriched environment can elicit NPAS4 expression in behaviorally-relevant neuronal populations throughout the hippocampus^6,13,22–25^. In CA1 pyramidal neurons, NPAS4 deletion highlights its central role in modulating inhibitory input: knockout neurons exhibit reduced somatic and increased dendritic inhibition compared to neighboring wild-type neurons^6^. Importantly, these changes arise from the selective regulation of synapses formed by anatomically distinct populations of cholecystokinin (CCK)-expressing interneurons^22,25,26^, indicating that NPAS4 expression reshapes how CA1 pyramidal neurons are integrated into the CCK inhibitory microcircuit.

During active exploration, a subset of CA1 pyramidal neurons fire in spatially selective patterns and show temporally organized spiking activity that is aligned to the ongoing theta rhythm. Individual pyramidal neurons can code for an animal’s location with spatially-tuned firing that occurs when the animal is in the neuron’s place field^27–30^. Place field firing is temporally coordinated with theta oscillations, which is the main oscillatory pattern during running. At the entrance into a place field, neurons fire late in the theta cycle, and at the exit, they fire early in the cycle. This phase precession orders the spiking of overlapping place fields such that a series of place fields along a spatial trajectory is also found in a time-compressed form within a theta cycle^31–37^. Recently, CCK+ inhibitory neurons have been shown to influence both the spatial and temporal firing of CA1 pyramidal neurons. Specifically, chronic dysregulation of CCK+ inhibitory neuron connectivity or deletion of cannabinoid receptors (CB1Rs) from CCK+ inhibitory neurons results in larger place fields recorded from CA1 pyramidal neurons^38,39^. Acute optogenetic silencing of CCK+ inhibitory neurons also leads to broader place fields and reveals a role for CCK+ inhibitory neurons in constraining burst firing and theta-phase precession of pyramidal neuron spiking^40^. Thus, the activity of CCK+ inhibitory neurons influences both the spatial and temporal tuning of pyramidal neurons, prompting us to hypothesize that NPAS4, as a transcriptional regulator of CCK+ inhibitory synapses, similarly refines pyramidal neuron firing.

To examine how NPAS4 influences spatial representations and temporal precision of spiking, we recorded CA1 pyramidal neuron firing in mice actively navigating a rectangular track, a behavioral context in which place fields and theta-phase–related firing patterns reliably emerge. We used viral expression of Cre to knockout *Npas4* and permit expression of Channelrhodopsin in the transduced CA1 pyramidal neurons. This enabled us to obtain extracellular optetrode recordings from intermingled wildtype (WT) and optotagged^41^ NPAS4 knockout (KO) neurons during navigation. Importantly, sparse deletion of NPAS4 allows for direct comparisons between neurons of different “genotypes” within an individual animal and reduces non-cell autonomous network dysregulation, such as seizure activity^5^, that occurs when NPAS4 is deleted from large populations of neurons. We observed that while NPAS4 KO neurons exhibited place fields, these fields were larger and less stable throughout the recording session in comparison to those recorded from WT neurons. In addition to these deficits in spatial representations, KO neuron firing was weakly coupled to theta oscillations and demonstrated less theta-phase precession. Collectively, these findings show that cell-specific loss of NPAS4 disrupts the spatial and temporal organization of CA1 pyramidal neuron activity, linking a transcriptional regulator of CCK+ inhibitory neurons to fundamental building blocks of higher-level cognition.

## RESULTS

### NPAS4 Reorganizes Somatodendritic Inhibition in CA1 Pyramidal Neurons of Adult Mice

We confirmed that in CA1 of adult mice, NPAS4 is expressed in select neurons following exposure to an enriched environment (EE; **Figure 1A**). We next asked whether NPAS4 has the same effect on inhibitory circuit organization in adult mice as has been previously shown in juveniles^6,22,26^. To test this, we injected AAV.CamKII_Cre-GFP into the CA1 region of adult *Npas4*^fl/fl^ mice (∼postnatal day 70, P70). After recovery from the surgery, mice were housed in EE for 2-3 months with regular updating of the environment to maintain novelty, matching the timeline used for separate experiments in which we obtained *in vivo* extracellular recordings. After the prolonged housing in EE, we prepared acute hippocampal slices and performed simultaneous whole-cell recordings from neighboring wild-type (WT) and NPAS4 knockout (KO) neurons. Electrical stimulation was delivered in the somatic or dendritic layers, and pharmacologically-isolated evoked inhibitory postsynaptic currents (eIPSCs) were recorded (**Figures 1B and C**). KO neurons had smaller amplitude somatic eIPSCs and larger amplitude dendritic eIPSCs than neighboring WT neurons (**Figure 1C**). Thus, NPAS4 shapes the distribution of inhibitory synaptic input onto CA1 pyramidal neurons in adulthood as it does in juveniles, and this synaptic phenotype persists with chronic exposure to EE.

**Figure 1.**
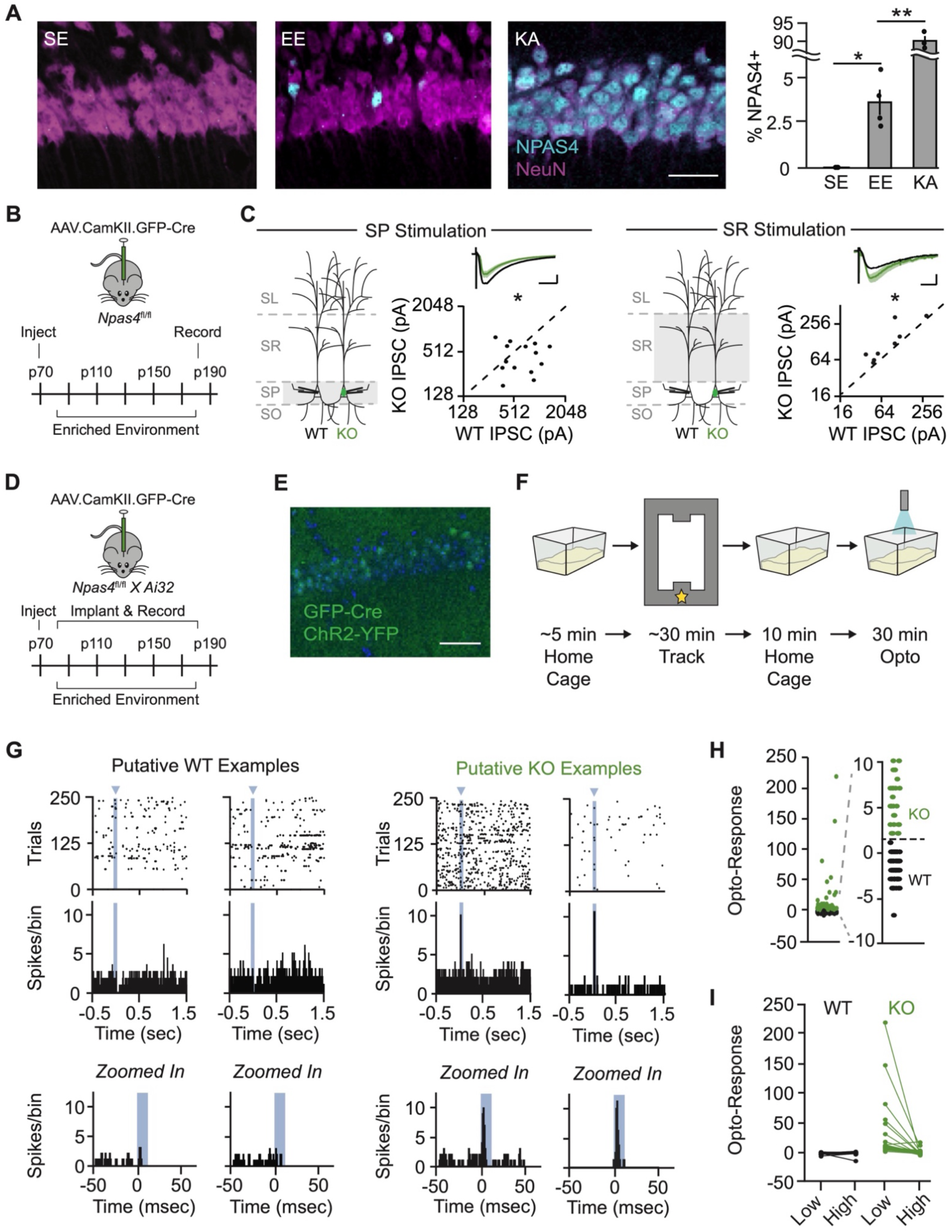
NPAS4 expression in CA1 pyramidal neurons in adult mice results in reorganization of inhibition along the somatodendritic axis. (A) NPAS4 expression in CA1 from mice housed in a standard environment (SE), an enriched environment (EE) for 90 minutes, or following kainic acid (KA) injection. Magenta: NeuN; teal: NPAS4. Scale bar = 50 µm. (Data are mean ± SEM; SE: N = 3 animals, 3 sections each; EE: N = 4 animals, 3 sections each; KA: N = 3 animals, 3 sections each; unpaired t test.) (B) Schematic of the experimental strategy and timeline for whole-cell patch-clamp recordings. (C) eIPSCs evoked in either stratum pyramidale (SP) or stratum radiatum (SR) recorded with simultaneous voltage-clamp from neighboring WT (black) and KO (green) pyramidal neurons. Geometric mean traces shown as percent of WT. SP scale bar = 20 ms, 25% of WT; SR scale bar = 20 ms, 50% of WT. (SP: N = 14 slices from 6 mice; SR: N = 8 slices from 5 mice; ratio paired t test.) (D) Schematic of the experimental strategy and timeline for *in vivo* electrophysiology recordings. (E) Example image of sparse KO used for in vivo recordings. Scale bar = 50 µm. (F) Schematic of the experimental timeline for each session. Star: reward location. (G) Example rasters (top) and peristimulus time histograms (PSTH; middle) from putative WT (left) and KO (right) neurons. (Bottom) Same PSTHs as above, shown at higher temporal resolution to highlight the 1-second window surrounding light stimulation. Blue arrow and bars: light stimulation (WT example 1: Animal 6 WT Cell 2; WT example 2: Animal 2 WT Cell 3; KO example 1: Animal 1 KO Cell 1; KO example 2: Animal 5 KO Cell 1). (H) Opto-response during low-power light stimulation (∼0.3 mW) in cells categorized as WT (black) or KO (green). Opto-response is defined as the peak PSTH value during light-on minus the peak PSTH value during light-off periods of the optostim protocol. (I) Opto-response during high-power light stimulation (>0.3 mW) in cells categorized as WT (black) or KO (green). As KO neurons were recruited into the pop-spike during high-power light stimulation, their opto-responses decrease substantially. *p < 0.05; **p < 0.01.

### Optical Tagging Enables *In Vivo* Identification of NPAS4 Knockout Neurons

We sought to determine how the loss of NPAS4 affects pyramidal neuronal firing in the context of an intact network with ongoing, behaviorally-driven network activity. To accomplish this, *Npas4^fl/fl^*mice were crossed to Cre-dependent Channelrhodopsin-2 (ChR2) mice (*Npas4^fl/fl^*:Ai32). Double-homozygous offspring were transduced with AAV.CamKII_Cre-GFP, resulting in an intermingled population of WT and KO neurons, where the KO neurons also expressed ChR2 (transduced neurons: 30-60%; **Figure 1D, E, and S1**). After recovery from surgery, mice were housed in EE for 2-3 months, including while electrophysiological recordings were obtained. Daily extracellular recordings were acquired first in the home cage, then during exploration of the rectangular track, and again in the home cage, with light stimulation delivered at the end of the final home cage session (**Figure 1F**). Spikes were sorted and clearly separated into clusters defined as a “unit”, likely corresponding to spikes generated by a single neuron. In addition, units were well-isolated in both the pre-track and post-track home cage recording periods, demonstrating physical stability of the tetrode across the recording session (N = 174 units from 8 male mice, **Figure S2E**).

During opto-recording periods (10-30 minutes) at the end of each session, light was delivered through the optical fiber of the optetrode (pulsed at 0.5 Hz; pulse duration of 10 msec) with the laser power set to evoke a small light-triggered response in the LFP without evoking a population spike (typically ∼3 mW; **Figures S2A and S2C**). Peristimulus time histograms (PSTH) were generated for each unit and the opto-response was calculated as the difference between the maximum spike count during the light pulse and the maximum spike count outside the pulse. Units that produced a reliable response to light stimulation were classified as NPAS4 KO neurons (opto-response > 1; n = 47 units), and units that did not were classified as “WT neurons” (opto-response < 1; n = 112 units; **Figure 1G-H**). Ambiguous units were excluded from further analysis (n = 15 units). KO neurons that express low levels of ChR2 might not robustly respond to the low-power light delivered, leading to some KO neurons being misclassified as WTs. To confirm the veracity of our WT population, we delivered high-power light, causing opto-triggered spikes from KO neurons to be recruited into the population spike and leaving WT neurons unchanged (**Figures 1I, S2B, and S2D**). Clusters of spikes from WT or KO neurons had comparable separation metrics and were present throughout the recording session (**Figure S2E-H**). Finally, percentages of KO neurons mirrored *post hoc* quantification of transduction density across animals (**Figures S2I and S2J**). Collectively, these analyses gave us confidence in our assignment of WT or KO neurons from tetrode recordings.

### Spatial Tuning Is Impaired in NPAS4 Knockout CA1 Neurons

Given the role of CCK+ interneurons in sculpting spatial tuning, we hypothesized that the dysregulation of inhibitory input onto NPAS4 KO neurons would lead to aberrant activity patterns during navigation. To explore this, we compared WT and KO neuron firing while mice ran laps on the track for a food reward. Each track recording session consisted of 10 trials run in one direction (an epoch) followed by 10 trials run in the opposite direction, alternating for 8 epochs or 30 minutes, whichever came first (**Figure 2A**). Trained behavior across the track was stereotyped, as assessed by velocity (**Figures 2B and S2K-M**). Across all trials on the track and considering only periods of time when the animal was running (velocity > 2 cm/sec), KO neurons had slightly but significantly higher firing rates than WT neurons (**Figure 2C**) but fewer spikes that occurred in bursts (defined as ISIs < 10 msec; **Figures 2D and 2E**).

**Figure 2.**
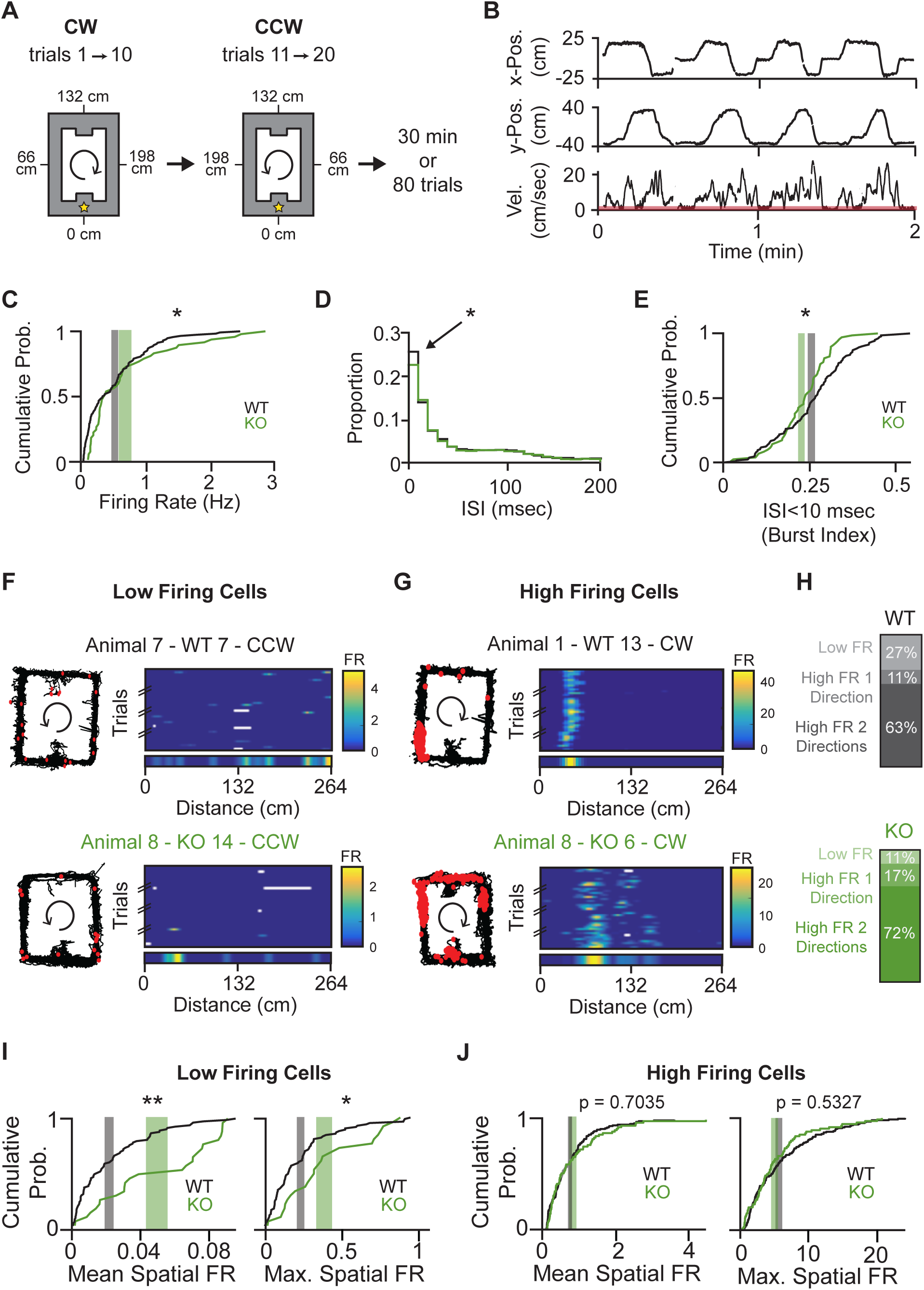
NPAS4 knockout neurons have fewer spikes in bursts and are more likely to have high firing rates, but high firing knockout and wild-type neurons exhibit comparable spatial firing. (A) Schematic of the track recordings. Mice ran in one direction on a rectangular track for 10 trials before being required to switch directions. This behavior was repeated for up to 80 trials or 30 minutes, whichever occurred first. The track was linearized as indicated. (B) Example traces showing x-position (top), y-position (middle), and velocity (bottom) as an animal ran along the track. Time periods during which velocity fell below 2 cm/sec (red bar on velocity plot) were excluded from analysis. (C) Cumulative probability distribution of firing rates across the session. Firing rate was defined as the total number of spikes divided by session duration. Gray shaded region: ± SEM for WT, centered at the mean; green shaded region: ± SEM for KO, centered at the mean. (WT: N = 112 cells; KO: N = 47 cells; Kolmogorov–Smirnov test.) (D) Histogram of interspike intervals (ISIs). Arrow indicates ISIs < 10 ms. (WT: N = 112; KO: N = 47; Kolmogorov–Smirnov test.) (E) Cumulative probability distribution of the proportion of spikes in bursts, defined as ISIs < 10 ms. Shaded regions as in (C). (WT: N = 112; KO: N = 47; Kolmogorov–Smirnov test.) (F) Example linearized rate maps from a ‘low firing’ WT (top) and KO (bottom) neuron. Left: trajectory of the session (black), with spikes (red dots). Right: trial-by-trial linearized rate map for one direction, with the trial-averaged rate map shown below. (G) As in (F), but for ‘high firing’ WT and KO neurons. (H) Percentage of cells classified as ‘low firing’ or ‘high firing’ in one or both directions (p < 0.05; chi-square goodness-of-fit test) (I) Cumulative probability distributions of mean and maximum spatial firing rates for ‘low firing’ cells. Shaded regions as in (C) (WT: N = 86; KO: N = 26; Kolmogorov–Smirnov test). (J) As in (I), but for ‘high firing’ cells. (WT: N = 138; KO: N = 68; Kolmogorov–Smirnov test.) *p < 0.05; **p < 0.01.

It was unclear whether the increased firing rate measured in KO neurons reflected a more robust response within a neuron’s place field or spurious firing outside the neuron’s spatial receptive field. To disambiguate this, we analyzed spatial firing rates by linearizing the track (reward zone at 0) and calculating the spike rate within 4 cm bins (for example, see **Figures S3A and B**) when the animal was running (velocity > 2 cm/sec). Place fields are direction-selective^42,43^; thus, trials run in opposite directions (clockwise (CW) or counterclockwise (CCW)) were analyzed independently and considered as distinct data points.

To focus our analyses on neurons that are active during behavior, we classified neurons as “high firing” if their mean spatial firing rate exceeded 0.1 Hz and their trial-averaged maximum spatial firing rate exceeded 1 Hz (**Figure 2G**); neurons below this cutoff were classified as “low firing” (**Figure 2F**). Histograms of mean and maximum spatial firing rates revealed a continuous distribution in both genotypes (**Figure S3C and D**), underscoring that this classification is not categorical but instead provides a pragmatic cutoff to exclude cells with minimal spatially organized spiking.

NPAS4 WT and KO neurons had dramatically different percentages of neurons with low and high spatial firing rates during behavior. Among the WTs, ∼27% of neurons had low firing rates in both directions, and 11% had low firing rates in one direction. In comparison, KO neurons were significantly different, with low firing rates in 11% of neurons in both directions and 17% in one direction (**Figure 2H**). Moreover, among the low firing rate neurons, KO neurons had significantly higher mean and maximum spatial firing rates (**Figure 2I**). In contrast, and despite the larger percentage of KO neurons in the high firing rate subgroup, we did not measure differences in the mean or maximum spatial firing rates between genotypes (**Figure 2J**). These results suggest that a greater proportion of KO neurons are active during behavior, possibly reducing the sparsity of the overall ensemble representation of the environment. Moreover, among the high firing rate neurons, the similarity between WT and KO spatial firing rates when averaged across the entire track raises the question of whether WT and KO neurons have comparable place field representations.

We considered the high firing rate neurons as putative place cells^29^ and defined a place field as the contiguous bins in which the firing rate was above 10% max firing and at least one bin was above 50% max (**Figure 3A**). Both WT and KO place cells had comparable numbers of place fields (typically only one; **Figure 3B**) that tiled the track and were directionally selective (**Figure 3C**) with low Pearson’s correlation coefficients between CW and CCW directions (**Figure S3E**). Despite these similarities, KO neurons generated significantly larger place fields than WT neurons (50 ± 3.52 cm and 39 ± 1.86 cm, respectively; **Figure 3D**).

**Figure 3.**
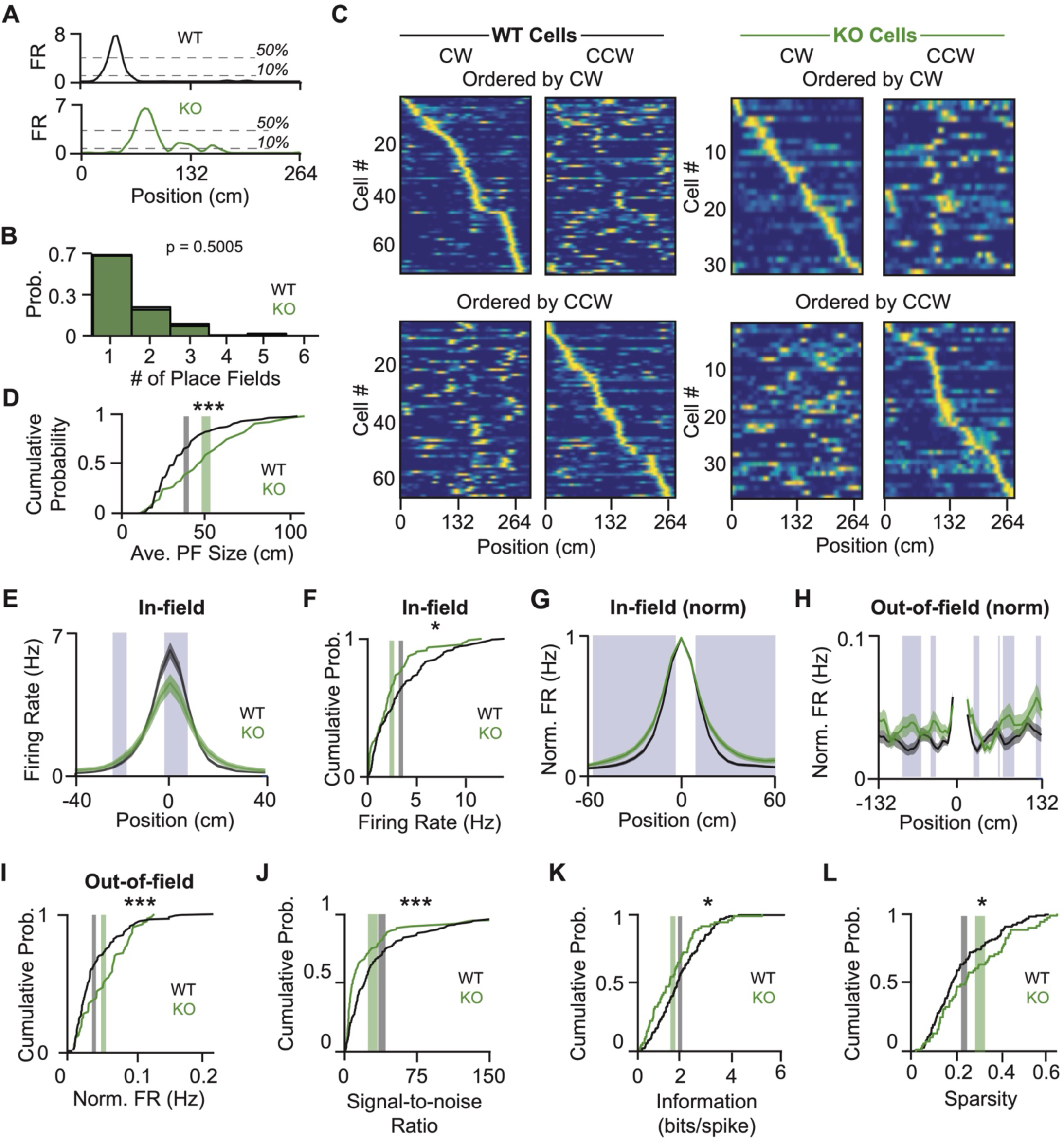
NPAS4 knockout neurons are less spatially tuned than simultaneously recorded wild-type counterparts. (A) Example trial-averaged rate maps from a WT (top) and KO (bottom) neuron. Dotted lines mark 10% and 50% of the peak rate and were used to identify place fields (WT example: Animal 1, WT Cell 35; KO example: Animal 6, KO Cell 2). (B) Histogram of the number of place fields per neuron (WT: N = 138; KO: N = 68; Mann–Whitney test). (C) Trial-averaged rate maps from WT (left) and KO (right) neurons. Top: ordered by the peak location in the clockwise (CW) direction, with the same order applied to activity in the counterclockwise (CCW) direction. Bottom: ordered by peak location in the CCW direction (WT: N = 138; KO: N = 68). (D) Cumulative probability distribution of average place field size. Place fields were defined as sets of contiguous bins above 10% of the peak that also included at least one bin above 50%. Gray shaded region: ± SEM for WT, centered at the mean; green shaded region: ± SEM for KO, centered at the mean (WT: N = 138; KO: N = 68; Kolmogorov–Smirnov test). (E) Trial-averaged in-field rates for WT and KO place fields, centered on the peak of each field. Blue shaded regions indicate bins with p < 0.05 (WT: N = 183 fields over 138 neurons; KO: N = 95 fields over 68 neurons; Kolmogorov–Smirnov test). (F) Cumulative probability distribution of average in-field firing rates. Shaded regions as in (D) (WT: N = 138; KO: N = 68; Kolmogorov–Smirnov test). (G) As in (E), but firing rates normalized to each neuron’s peak. (H) As in (G), but showing only out-of-field bins. (I) Cumulative probability distribution of average out-of-field firing rates. Shaded regions as in (D) (WT: N = 138; KO: N = 68; Kolmogorov–Smirnov test). (J) Cumulative probability distribution of signal-to-noise ratio, defined as the average in-field firing rate divided by the average out-of-field firing rate per neuron. Shaded regions as in (D) (WT: N = 138; KO: N = 68; Kolmogorov–Smirnov test). (K) Cumulative probability distribution of spatial information. Shaded regions as in (D) (WT: N = 138; KO: N = 68; Kolmogorov–Smirnov test). (L) Cumulative probability distribution of sparsity. Shaded regions as in (D) (WT: N = 138; KO: N = 68; Kolmogorov–Smirnov test). *p < 0.05; ***p < 0.001.

Our observation that WT and KO neurons have comparable spatial firing rates across the entire track, but KO neurons have larger place fields and thus larger portions of the track with elevated firing, presents a conundrum. One possible way to reconcile these measurements, particularly considering the excessive dendritic and reduced somatic inhibition received by NPAS4 KO neurons, would be if KO neurons have relatively lower firing rates within the place field and higher firing rates outside of the place field. Among WT neurons, 69% of action potentials were within the place field (”in-field”) and 31% were distributed across the rest of the track (“out-of-field”); for KO neurons these values were 63% and 37%, respectively (**Figure S3F**). To quantify in-field spatial firing rates, we aligned each field to the peak and compared the firing rates in each spatial bin (**Figure 3E**). Across the population, KO neurons had significantly lower peak firing rates and reduced in-field firing rates (**Figures 3E, 3F**). After normalizing to the peak firing, KO neurons showed significantly higher firing rates as the animal entered and exited the field, driving the larger normalized place fields (**Figure 3G**). Moreover, increasing the firing rate threshold for defining a place field eliminated the differences between genotypes (**Figures S3G and S3H**), suggesting that place fields generated by KO neurons are not scaled versions of WT place fields.

If the overall spatial firing rates are comparable but the in-field firing rate is lower, KO neurons must have higher out-of-field firing. Indeed, in our normalized, peak-aligned rate maps, KO neurons often had higher out-of-field firing rates, and this was significant when averaging across all out-of-field bins (**Figures 3H, 3I**). The shift in spikes from in-field (“signal”) to out-of-field (“noise”) strongly reduced the KO neurons’ signal-to-noise ratio (**Figure 3J**). Indeed, KO place cells conveyed less spatial information with each spike and fired more uniformly over the track (**Figures 3K, 3L**). These results were robust to variations in firing rate threshold (**Figures S4A-C**) and persisted when controlling for firing rate (**Figures S4D-G**). Thus, deleting NPAS4 reduces the precision of CA1 pyramidal neuron spatial representations.

### Stability of Spatial Firing Is Reduced in NPAS4 Knockout Neurons

In the average rate maps, the KO neurons have significantly broader spatial tuning than WT neurons. Surprisingly, this difference seemed less prominent when looking at individual trials (**Figures S5A–F**). Place fields that shift location on a trial-to-trial basis could give the appearance of a larger average field while masking an underlying instability of the field. To assess place field stability, we averaged firing across the track for each epoch (10 trials) and calculated Pearson’s correlation coefficients (PCCs) between successive epochs for WT and KO neurons. While both WT and KO neurons had PCCs that were higher than shuffles, the PCC was lower for KO neurons at each comparison, demonstrating that the overall spatial firing was less consistent across epochs (**Figures 4A, 4B, and S5G**). Restricting the analysis to in-field or out-of-field bins also revealed lower correlations in KO neuron firing (**Figure S5H and S5I**), suggesting that their place fields are more labile, and that the higher out-of-field activity measured in the KO neurons is spurious and unlikely to be nascent field formation.

**Figure 4.**
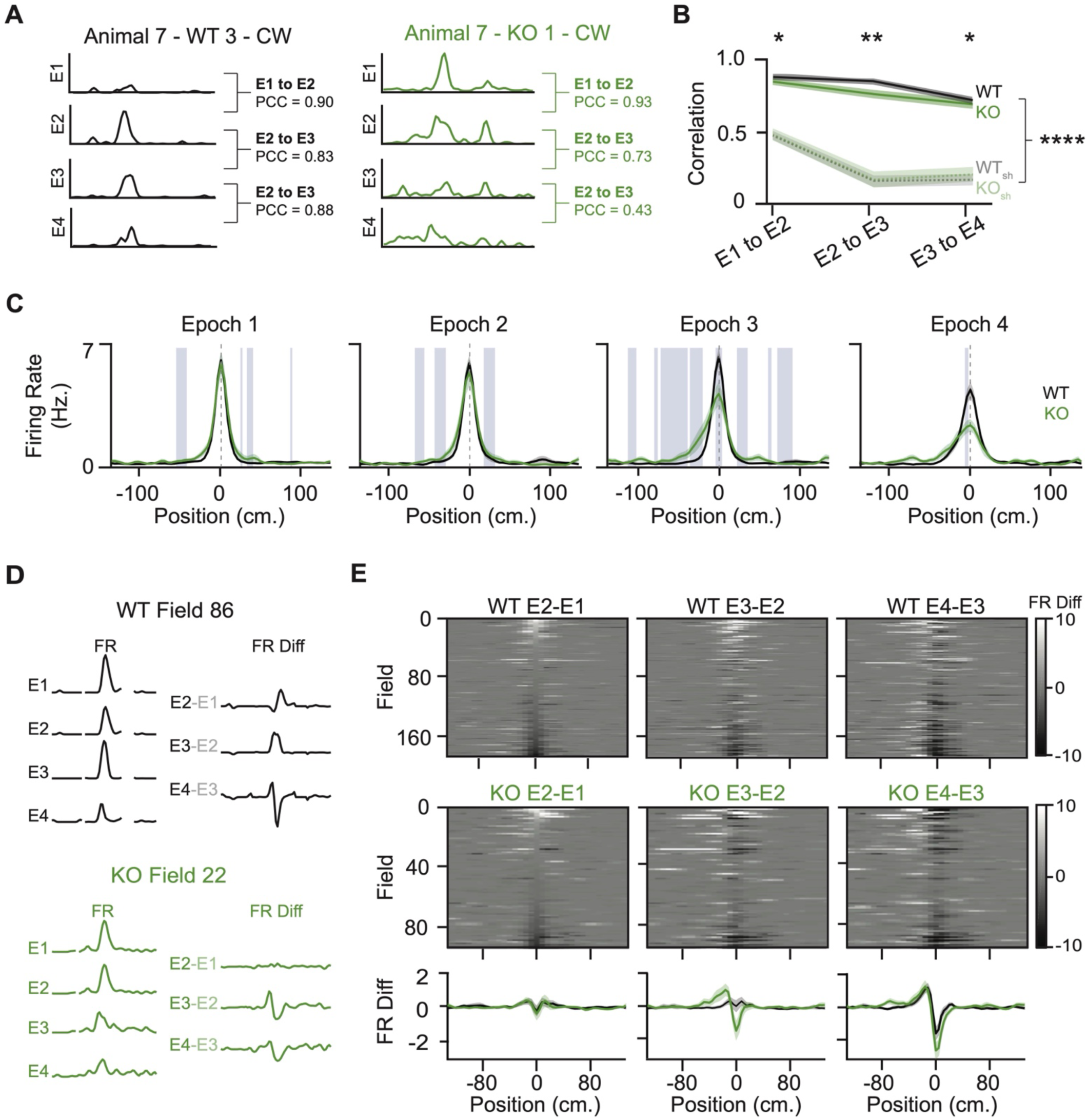
NPAS4 knockout neurons are less stable and shift their place fields towards the field entrance more rapidly than wild types. (A) Example trial-averaged rate maps from each of the four epochs (sets of 10 trials) for a WT (left) and KO (right) neuron. For each pairwise comparison (epoch 1 to 2, epoch 2 to 3, and epoch 3 to 4), the Pearson’s Correlation Coefficient (PCC) was calculated. (B) PCC across sequential epoch comparisons for WT (solid black) and KO (solid green) neurons, alongside shuffled controls (WT shuffle: dotted gray; KO shuffle: dotted green). Shuffled distributions were generated by spatially shifting trials randomly (WT: N = 138; KO: N = 68; Kolmogorov–Smirnov test for WT vs. KO; Wilcoxon signed-rank test for comparisons to shuffled distributions). (C) Trial-averaged rate maps for WT and KO place cells, separated by epoch and centered on the peak in Epoch 1. Blue shaded regions indicate bins where p < 0.05 (WT: N = 138; KO: N = 68; Kolmogorov–Smirnov test). (D) Example showing how difference maps are generated: trial-averaged rate maps from each epoch are centered on the peak of Epoch 1, and the difference between sequential epochs is computed. (E) Difference maps for WT (top) and KO (bottom) place fields across all epoch comparisons. Mean difference maps across neurons are shown below (WT: N = 176 fields from 138 neurons; KO: N = 91 fields from 68 neurons). *p < 0.05; **p < 0.01; ****p < 0.0001.

Is the reduced correlation across epochs due to a systematic change in KO neuron spatial firing? To assess this, each place field was aligned to its peak location in the first epoch (E1; **Figure 4C**) and fields from all neurons were averaged within each epoch. In WT neurons, place fields remained stable across the first three epochs, but in E4, they showed a reduction in peak firing and a broader tail extending toward the field entrance—hinting at the Mehta effect^44,45^ and behavioral timescale plasticity^46^. In contrast, KO neurons were indistinguishable from WTs in E1, but as early as E2, the average field began to degenerate, with reduced peak firing and increased activity in peri-field and out-of-field regions. These differences were exacerbated in the population averages in E3 and E4, indicating substantial heterogeneity in the changes exhibited by KO neurons.

To better understand how place fields changed at the level of individual neurons, we examined the shift in peak firing location across epochs. WT neurons showed minimal movement of the place field peak through E3, but by E4, over half of the fields had shifted toward the field entrance. KO neurons also showed shifts in peak firing towards the entrance, but these differences emerged as early as E2 (**Figure S5J and S5K**). To visualize changes in spatial firing across the entire track—not just at the peak—for each neuron and across epochs, we computed firing rate difference maps between successive epochs. Each map was centered on the peak firing location from E1, and fields were ordered based on the change in firing between E2 and E1 across all comparisons (**Figures 4D and 4E**). Although there were a variety of responses within both populations, many place fields from WT neurons showed the characteristic Mehta effect at the level of individual cells, most consistently between E3 and E4^44,45^. KO neuron place field locations were more variable, with individual field positions jumping larger distances, often in early epochs, and without stability of the new location in subsequent epochs. Collectively, these analyses show that while place fields from both WT and KO neurons shift over time, the KO neuron place fields are exceptionally labile, likely underlying the larger trial-averaged field sizes and emerging from the dysregulation of inhibition.

### Temporal Precision of Spiking Is Impaired in NPAS4 Knockout CA1 Neuron

During running, CA1 pyramidal neuron firing is organized with respect to the underlying theta rhythm in the local field potential (LFP). While phase precession broadens the range of theta phases at which CA1 pyramidal neurons fire, spiking remains largely confined to a preferred portion of the theta cycle^31–34—a^ constraint shaped in part by rhythmic inhibition^40,47–50^. This led us to examine whether theta-phase coupling is disrupted in KO neurons.

In our experiments, KO and WT neurons are intermingled; hence, spikes generated by neurons of either genotype are aligned to a shared LFP. During bouts of running, the theta power in the LFP was indistinguishable from control-injected animals, indicating that sparse deletion of *Npas4* did not impact theta oscillations (**Figure S6A and B**). We filtered the LFP (2-20 Hz) to focus on the theta band and obtained the phase of theta at which each spike occurred, where 0° is the peak of theta and 180° is the trough (**Figure 5A**). We fit a vector to the spike-theta phases and used this to calculate the mean vector length (MVL) and the preferred theta phase for the best field (field with the highest firing rate) of each cell (**Figure 5B**). Although we did not observe any significant differences in the preferred theta phase between WT and KO populations (**Figure 5C and D**), we did find that KO cells had significantly lower MVLs (**Figure 5E**), indicating that the KO neurons have a weaker phase preference. This difference persisted when we considered in-field activity from all fields. No difference between WT and KO theta-phase coupling was detected when restricting the analysis to out-of-field firing **(Figure S6C-E)**. The strength of theta coupling varies as an animal traverses a given neuron’s place field, decreasing at the peak of the field^51,52^. We observed this pattern for both WT and KO firing. KO neurons, however, had considerably lower MVLs regardless of place field position (**Figure 5F**). Taken together, these data demonstrate that KO neuron place field firing is less theta-coupled than WT neuron counterparts.

**Figure 5.**
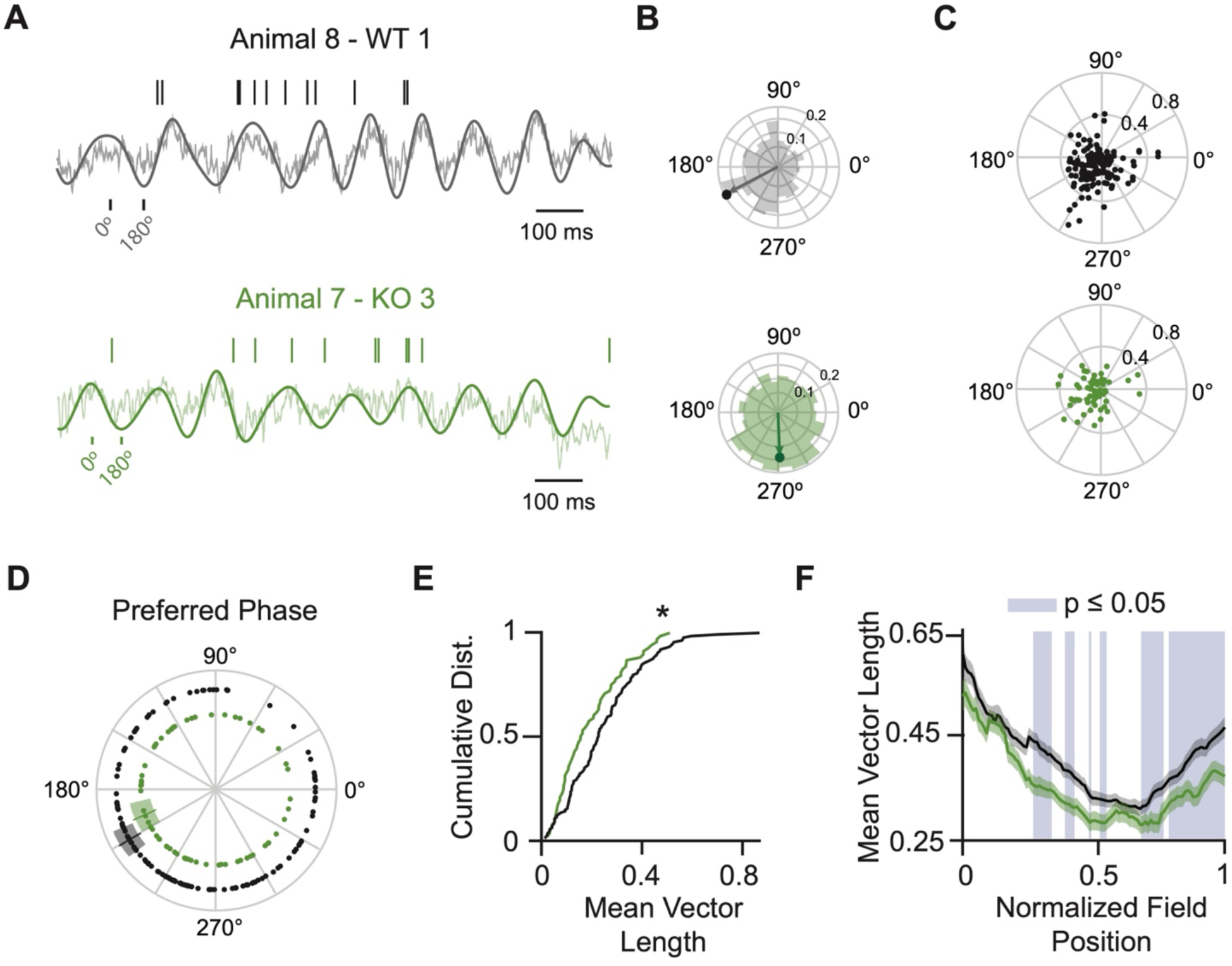
NPAS4 knockout cells are less theta-coupled than wild-type counterparts. (A) Example of spiking relative to the local field potential (LFP) for WT (top) and KO (bottom) neurons. Spikes are shown as vertical lines above the raw LFP trace with the theta-filtered LFP overlaid. (B) Rose plots show the theta phase of all spikes from the same example neurons with the mean vector overlaid. (C) Rose plot scatter for all neurons. Angular position indicates preferred theta phase; radial position indicates mean vector length (WT: N = 138; KO: N = 68). (D) Preferred phase for WT and KO neurons. Solid line: mean; shaded region: SEM. Concentric circles shown for visualization only (WT: N = 138; KO: N = 68). (E) Cumulative probability distribution of mean vector lengths. Gray shaded region: ± SEM for WT, centered at the mean; green shaded region: ± SEM for KO, centered at the mean (WT: N = 138; KO: N = 68; Kolmogorov–Smirnov test). (F) Mean vector length as a function of normalized field position. Blue shaded regions indicate bins where p < 0.05 (WT: N = 183 fields from 138 neurons; KO: N = 95 fields from 68 neurons; Kolmogorov–Smirnov test). *p < 0.05.

Phase precession relies on spike timing across the theta cycle, wherein a pyramidal neuron’s spikes occur at earlier phases of theta as the animal moves through its place field. This enables information about the sequential activation of fields to be preserved in the temporal ordering of spikes for fields with spatial overlap. Since KO neurons have reduced theta-phase coupling, we extended our analysis to phase precession to determine if there are also differences. For each trial, we plotted each spike’s theta phase relative to the animal’s location in the place field (**Figure 6A**) and used the slope of the linear fit to quantify phase precession (**Figure 6B and C**)^37,53,54^. For trials that met established criteria^55^ (**Figure S6F**), trial-to-trial variability in slope did not differ between genotypes (**Figure S6G**). However, the mean slope per neuron (**Figure 6D**; median slopes: **Figure S6N**)—a measure of overall phase precession strength—was significantly shallower in KO neurons, despite both groups showing the expected negative values indicative of phase precession. This difference disappeared when we shuffled spike phases or spatial positions (**Figure S6H-K**) and remained robust to bootstrapping (**Figures S6L and S6M**). Thus, NPAS4 KO neurons have weaker phase precession than WT neurons.

**Figure 6.**
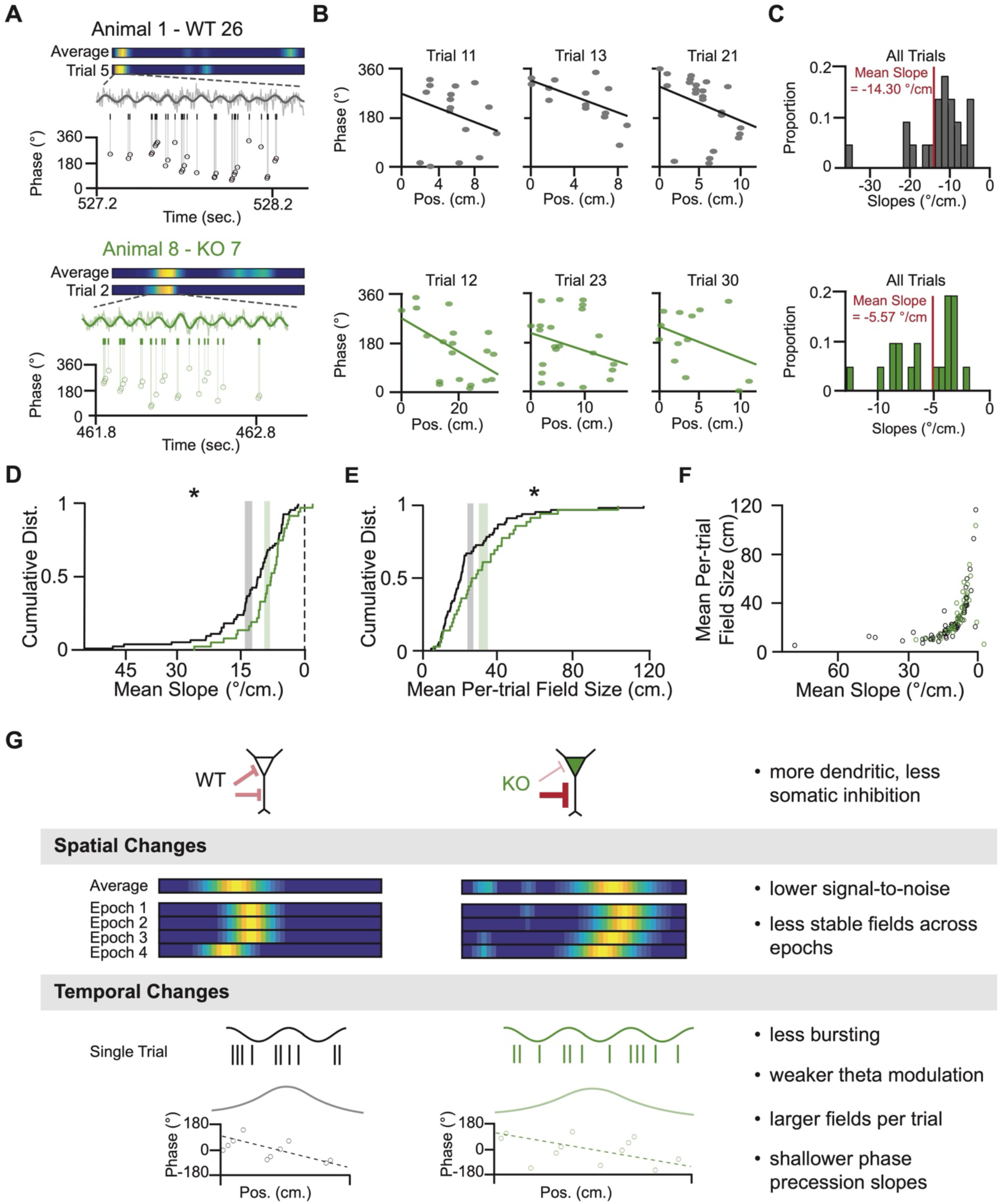
NPAS4 KO neurons exhibit shallower phase precession slopes compared to WT counterparts. (A) Example WT (top) and KO (bottom) neurons showing theta-related spiking during a single field pass. For each example, the top panel shows the trial-averaged rate map across all trials; the panel immediately below shows the rate map for one example trial. The raw local field potential (LFP) during that trial is shown with the theta-filtered LFP overlaid. Spikes are marked as vertical lines below the LFP, and the theta phase of each spike is plotted over time. (B) Three example trials from the neurons shown in (A). Scatter plots show theta phase of each spike over position with linear fits overlaid. (C) Histogram of phase precession slopes across all trials for the neurons in (A). Red line: median slope. (D) Cumulative probability distribution of mean phase precession slopes. Only the field with the highest firing rate from each neuron is included. Gray shaded region: ± SEM for WT, centered at the mean; green shaded region: ± SEM for KO, centered at the mean (WT: N = 70; KO: N = 36; Kolmogorov–Smirnov test). (E) Cumulative probability distribution of mean field size calculated on a trial-by-trial basis. Only cells included in (D) are included. Gray shaded region: ± SEM for WT, centered at the mean; green shaded region: ± SEM for KO, centered at the mean (WT: N = 70; KO: N = 36; Kolmogorov–Smirnov test). (F) Mean field size of each neuron plotted against its mean phase precession slope. Black: WT; Green: KO (WT: N = 70; KO: N = 36; Kolmogorov–Smirnov test). (G) Schematic summarizing results. *p < 0.05.

Larger place fields have shallower phase precession^54,56^, raising the possibility that field size differences could account for the diminished phase precession observed in KO neurons. First, considering the trials that met the criteria above and calculating field size for each trial independently, we compared the average per-trial field sizes between the WT and KO neurons. KO neurons had significantly larger per-trial field sizes than WTs (**Figure 6E**), although both were smaller than those calculated using the complete set of trials (**Figure S5D**). This result is consistent with the idea that our phase precession criteria selected for neurons with stronger spatial tuning. For both WT and KO neurons, field size and phase precession slope were highly correlated (**Figure 6F**) (Spearman’s correlation: WT: Rho = 0.7754, p = 1x10^-10^; KO: Rho = 0.7650, p = 1x10^-10^). To determine the relative contributions of genotype, field size, and theta modulation strength to the difference in phase precession, we built a linear regression model, using these three variables as predictors (**Table S1**); field size and slope were log transformed to account for the non-linearities in these variables (**Figure S6O and P**). The model explained 28% of the variance in slope (adjusted R² = 0.2784) with field size emerging as the only significant positive predictor of phase precession (β = 0.9191, p = 1.3433 x 10^-12^). Thus, the reduction in phase precession observed in NPAS4 KO neurons is linked to the concomitant increase in place field size, suggesting a coupling between spatial representations and temporal precision of spiking that is perturbed when NPAS4-dependent regulation of inhibition is disrupted.

## DISCUSSION

NPAS4 orchestrates a reorganization of inhibitory inputs along the somatodendritic axis of CA1 pyramidal neurons, serving as a molecular link between a neuron’s recent activity and targeted changes in inhibitory synapse composition. Here, we demonstrate that NPAS4-mediated reorganization of inhibition occurs in adult animals, not just in juveniles, suggesting that NPAS4 is important for tuning neuronal activity across the lifespan of the animal. Using a sparse knockout strategy and *in vivo* optotagging to differentiate between WT and KO neurons within awake, behaving animals, we identified several consequences of the loss of NPAS4, schematized in **Figure 6G**. First, we showed that NPAS4 KO neurons have more uniform firing across the track. Specifically, KO neurons exhibit lower firing rates within the neuron’s place field and higher firing rates across the rest of the track. In addition to the degenerate representation of space, KO neuron place fields were less stable, shifting large distances over few trials. The temporal organization of spikes was also disrupted—KO neurons showed weaker theta coupling and exhibited shallower phase precession. Together, these findings reveal that NPAS4 expression has substantial and lasting consequences for the spatial precision and temporal organization of CA1 pyramidal neuron firing in adult mice.

Our experimental design provides insight into the cell-autonomous effects of behavioral experience on pyramidal neuron activity. By sparsely knocking out NPAS4 in adult mice, we reduce the likelihood of developmental or circuit-level compensations that could mask or exaggerate NPAS4-dependent phenotypes. While our manipulation is sparse, NPAS4 is knocked out for at least one month. Additional methodological innovation is needed to examine the more acute consequences of NPAS4 expression or deletion. Our use of electrophysiological recordings from intermingled WT and KO neurons enables precise measurement of spike timing relative to a shared local field potential (LFP)—a level of temporal resolution not achievable with calcium imaging^57,58^. This study advances our understanding of how an activity-dependent transcription factor shapes *in vivo* information encoding and creates a bridge between molecular and circuit-level biology.

Detailed knowledge of how NPAS4 changes inhibitory synaptic connectivity is an essential context for understanding the results in this study and reveals important future questions to be explored. NPAS4-dependent changes in CCK+ inhibition are necessary for the emergence of dendritic plasticity mechanisms^26^. We speculate that the increased dendritic inhibition observed in NPAS4 KO neurons suppresses burst firing and impairs dendritic plasticity—processes thought to stabilize place fields^46,59–69^. In contrast, reduced somatic inhibition may permit spikes to be generated when they should not be, leading to spurious out-of-field firing. Inhibition from CCK+ neurons onto CA1 pyramidal neurons has been shown to constrain place field size and increase stability^38^. Our findings support this role and extend it by linking activity-dependent expression of NPAS4 to the same spatial receptive field properties.

Beyond their role in shaping pyramidal neuron spatial receptive fields, CCK+ inhibitory neurons also contribute to the temporal organization of spiking through their influence on theta-modulated firing. *In vivo* recordings obtained from anesthetized rats^50^ and awake behaving mice^40^ have shown that CCK+ basket cell activity *itself* is theta-modulated, though the preferred phase of firing varies between paradigms. In principle^50,70^, the rhythmicity of CCK+ basket cell firing imposes a window of opportunity for pyramidal neuron firing. Reduced CCK+ basket cell inhibition, as demonstrated in NPAS4 KO neurons^22^, likely broadens this window thereby contributing to the diminished theta coupling observed in this study. While CCK+ inhibitory neurons have not been directly linked to phase precession, activation of CB1 receptors (found exclusively on CCK+ inhibitory neurons in the hippocampus) has been shown to disrupt phase precession^71,72^. We observe a clear disruption of this temporal coding property in NPAS4 KO neurons. Notably, phase precession slope was significantly correlated with place field size in our dataset—a relationship that also emerged as a key predictor in our regression model. Although others have reported similar correlations, the mechanistic link between field size and precession slope remains unresolved: does one drive the other, or do both arise from a shared upstream process? A major challenge for the field will be to develop strategies that disentangle these interdependent coding features and clarify how distinct forms of inhibition interact to shape them. Nevertheless, our findings demonstrate that the distribution of place field sizes and phase precession slopes are shifted in NPAS4 KO neurons, suggesting a shared dependence on NPAS4-mediated inhibitory synapse organization.

The activity of CCK+ inhibitory neurons is strongly modulated by behavioral state, suggesting that their influence on pyramidal neuron output may be particularly important during transitions in network dynamics. For example, CCK+ inhibitory neurons increase their firing during shifts from locomotion to immobility^73,74^, and are particularly recruited during hippocampal ripples in NREM sleep^40^. Based on this, NPAS4 may be instrumental in suppressing pyramidal neuron firing during periods of rest and preventing indiscriminate recruitment into ripple-delimited replay events. While our study focused on periods of active exploration, future work examining how NPAS4 shapes CA1 activity during transitions to immobility or during specific stages of sleep, is likely to be especially relevant.

Finally, there are many activity-dependent transcription factors, in addition to NPAS4, that are expressed in neurons. Single-cell transcriptomics show us that the expression of these transcription factors often overlaps. For example, NPAS4+ neurons are often also Fos+. Yet, these two transcription factors drive different mutually-exclusive synaptic phenotypes, regulating somatic inhibition from CCK+ or PV+ basket cells, respectively^22,75^. Moreover, there is recent evidence that Fos also shapes spatial representations in CA1 pyramidal neurons, with high Fos expression correlating with larger, more stable place fields^16^. To understand the rich complexity of activity-dependent gene regulation, future studies that closely examine the synergistic, antagonistic, or mutually exclusive actions of different activity-dependent transcription factors are needed.

## Resource availability

### Lead contact

Requests for further information, resources, and reagents should be directed to and will be fulfilled by the lead contact, Brenda Bloodgood.

### Materials availability

This study did not generate new, unique reagents.

### Data and code availability

The electrophysiology data will be deposited on Zenodo upon publication of the study. Code is available at https://github.com/Bloodgood-Lab/Payne-et-al/tree/main.

## Acknowledgments

We thank members of the Bloodgood lab for ongoing discussion and input that shaped the direction of this study. We specifically acknowledge Kayla Torres, Destiny Tellez, Jedd Santamaria, Hunter Robbins, Jacob Gerzenshtein, Jerry Hou, Elena Dreisbach, and Paola Guerrero-Servin for assistance with training mice and animal care. We are grateful to all the members of the Leutgebs’ lab for thoughtful discussion throughout the development and execution of this study. We specifically thank Sunandha Srikanth, Ipshita Zutshi, Geoff Diehl, Sia Ahmadi, and Li Yuan for their thoughtful discussions and assistance with code used for analyses. The authors acknowledge with gratitude Nicholas Topper, Sarah Burke, and Andrew Maurer for developing the synchronization technique used in this study to align electrophysiology and video recordings. This work was supported by the National Institutes of Health through grants to B.L.B. (NINDS R01 NS111162) and training support to A.P. (NIMH F31 MH123112, NIBIB T32 EB009380, and NINDS T32 NS061847).

## Author Contributions

Conceptualization, A.P, A.L.H, and B.L.B.; methodology, A.P., C.S., and C.Q.; formal analysis, A.P., C.S., and D.A.H.; investigation, A.P., D.A.H., L.L.H., and R.S.G.; resources, J.K.L., S.L., and B.L.B.; writing - original draft, A.P. and B.L.B.; writing - review and editing, A.P., D.A.H., A.L.H., J.K.L., S.L., and B.L.B.; visualization - A.P.; supervision, J.K.L., S.L., and B.L.B.; funding acquisition – A.P., and B.L.B.

## METHODS

### Mice

All experiments were conducted in accordance with National Institutes of Health (NIH) guidelines and following the approval of our protocol by UC San Diego’s Institutional Animal Care and Use Committee (IACUC). An *Npas4*^fl/fl5^ animal line was used for acute slice electrophysiology experiments and an *Npas4*^fl/fl^:Ai32 (Ai32 RRID:IMSR_JAX:012569) animal line was used for NPAS4 immunohistochemistry (IHC) and all *in vivo* electrophysiology experiments. Only male mice were used for the sparse *in vivo* experiments. All electrophysiology experiments were performed on adult animals (P70-P200) that were housed long-term in enriched environments. The enriched environment consisted of a running wheel, toys, wooden blocks, and other objects of various shapes, colors, and textures. To maintain novelty, toys were replaced every two days. For all electrophysiology experiments, mice were kept in the vivarium on a reverse 12-hour light-dark cycle and were single-housed.

### NPAS4 Immunohistochemistry

For NPAS4 staining in enriched animals, adult mice (P70-P200) housed in a normal light-dark cycle were removed from the vivarium and left in a dark, empty room for two hours prior to the experiment. Half of the mice were then placed into an enriched environment (EE) for 90 minutes. In a separate set of experiments, additional mice received an intraperitoneal injection of kainic acid (KA; 2.5 mg/kg) and were placed in a large rat cage for 90 minutes. All KA-injected mice exhibited clear behavioral seizures during this period. The other half were left in their home cages for standard environment (SE) control. EE consisted of a large (2x2 ft) cardboard box with colorful patterns taped to the walls; two running wheels; plastic toys of various sizes, shapes, and colors; and wooden blocks. Mice were monitored for the full 90 minutes and continued to engage in the environment and actively explore for the majority of the enriched exposure. At the end of 90 minutes, mice from EE and SE were immediately anesthetized in isoflurane. The brains were extracted, the hippocampi dissected, and drop-fixed in 4% PFA for 2 hours. After 2 hours, the dissected hippocampi were rinsed in three ten-minute washes in 1X phosphate-buffered saline (PBS) before being moved to a 30% sucrose solution. The hippocampi were left in 30% sucrose overnight or until they had sunk to the bottom. The hippocampi were then frozen in OCT (optimal cutting temperature) compound and sectioned along the dorsoventral axis using a cryostat. Sections from the dorsal hippocampi were selected and blocked overnight in 10% goat serum/0.25% triton-X in PBS. Primary antibody solutions were applied to the slides and consisted of 1% goat serum, 0.25% triton-X, primary antibody against NPAS4 (1:500; RbαNPAS4; ^5^, and primary antibody against NeuN (1:1000; GPαNeuN; Synaptic Systems RRID:AB_2619988) in 1X PBS. The primary antibody solution was left for 48 hours at which point the slides were rinsed with three 10-minute washes of 1X PBS. Secondary antibody solutions were applied and consisted of 1% goat serum, 0.25% triton-X, Alexa 568 secondary antibody (1:1000; GtαRb; ThermoFisher; RRID: AB_10563566), and Alexa 647 secondary antibody (1:1000; GtαGP; ThermoFisher; RRID: AB_2535867) in 1X PBS. The secondary solution was left for 24 hours at which point the slides were rinsed with three 10-minute washes of 1X PBS. Slides were cover slipped using Fluoromount with DAPI and imaged at 60X using a confocal microscope. Images were acquired on an Olympus Fluoview 1000 confocal microscope (× 60/1.42 [oil] plan-apochromat objectives; UC San Diego School of Medicine Microscopy Core).

### Image Quantification

To quantify the number of neurons expressing NPAS4 in the CA1 pyramidal cell layer, we manually counted the number of cells somatically expressing NPAS4 and divided by the total number of cells in the pyramidal cell layer (NeuN+). We used the same levels for all images and only counted cells as NPAS4+ if we observed NPAS4 signal in at least 50% of the soma as identified using our NeuN signal.

### Viruses

For all sparse infection experiments (*ex vivo* and *in vivo* electrophysiology), an adeno-associated virus (AAV) expressing Cre-GFP was used (pENN.AAV.CamKII.HI.GFP-Cre.WPRE.SV40 AAV9; Addgene Item ID:105551-AAV9). To achieve a sparse infection, the virus was diluted 1:3 or 1:4 with sterile saline just before injection.

### AAV Injections

All surgeries were performed in accordance with (NIH) guidelines and following the approval of our protocol by UC San Diego’s IACUC. Stereotaxic viral injection surgeries were performed on adult animals (P70). Animals were injected with flunixin (2.5 mg/kg) subcutaneously pre-operatively and post-operatively every 12 hours for 72 hours. Animals were anesthetized with isoflurane for the duration of the surgery (1.5%-2% isoflurane vaporized in oxygen) and body temperature was maintained at 37° C using a delta phase pad. Following induction of anesthesia, the mice were placed in a stereotaxic apparatus, the fur covering the scalp was shaved cleanly, and the scalp was cleaned with three iterations of betadine and 70% ethanol. An incision along the midline was made to expose the skull so that bregma and lambda could be observed. For both *ex vivo* (targeting medial CA1) and *in vivo* (targeting dorsal CA1) electrophysiology experiments the distance between bregma and lambda was used to scale the anterior-posterior (AP) coordinates.

For subsequent *ex vivo* experiments two burr holes were drilled bilaterally (four in total) above medial (along the dorsoventral axis) CA1. AP coordinates were calculated using an equation derived from successfully targeted surgeries. The equations for the AP coordinates were AP = (-2.30/3.14)*bregma lambda distance and AP = (-2.60/3.14)*bregma lambda distance. The medial-lateral (ML) coordinates were ±3.30 mm and the dorsal-ventral (DV) coordinates (three injections per burr hole) were -1.40, -2.50, and -3.60. For subsequent experiments in freely behaving mice, one burr hole was drilled above dorsal CA1 in the right hemisphere only. The equations for the AP coordinates were AP = (-1.44/3.14)*bregma lambda distance. The ML coordinates were ML=1.45 to 1.55 mm and the DV coordinates were DV=1.45 to 1.55 mm.

At each injection site virus was injected (*ex vivo:* 150 nL at each injection site; freely behaving: 300 nL; 100 nL/min) using a Hamilton syringe attached to a Micro4 MicroSyringe Pump Controller (World Precision Instruments). Three minutes post-injection, the needle was retracted, the scalp sutured, and the mouse recovered at 37° C before being moved to a new home cage in which it was individually housed for the duration of the experiments.

### Acute Slice Preparation

Transverse hippocampal slices were prepared from *Npas4^fl/fl^* mice (P176-P184) at least three months after stereotaxic injection of AAV.Cre-GFP into CA1. Animals were anesthetized briefly by inhaled isoflurane and decapitated. Blocking cuts were made to isolate the portion of the cerebral hemispheres containing the hippocampus and slice preparation was prepared as described previously ^22^. Specifically, hemispheres were mounted on a Leica VT1000S vibratome and bathed in NMDG-HEPES recovery solution (NMDG 93 mM, HCl ∼93 mM, KCl 2.5 mM, NaH2PO4 1.2 mM, NaCO3 30mM, HEPES 20mM, glucose 13 mM, NAC 12mM, sodium ascorbate 5mM, thiourea 2mM, sodium pyruvate 3mM, MgSO4 10mM, CaCl2 0.5mM, 300-310 mOsm, pH 7.3-7.4 with HCl, saturated with 95% O2/5% CO2). After cutting, sections were transferred to 34° NMDG-HEPES recovery solution and sodium was spiked in over 30 minutes as previously described ^76^. Slices were then transferred to modified HEPES holding ACSF ^76^ (NaCl 92mM, KCl 2.5mM, NaH2PO4 1.2 mM, NaHCO3 30mM, HEPES 20mM, glucose 13 mM, NAC 12mM, sodium ascorbate 5mM, thiourea 2mM, sodium pyruvate 3mM, MgSO4 2mM, CaCl2 2mM, 300-310 mOsm, pH 7.3-7.4 with NaOH, saturated with 95% O2/5% CO2) where they were recovered for 1 hour and then maintained for the remainder of the day (∼6 hr).

### *Ex Vivo* Electrophysiology and Pharmacology

Infection density varied with distance from the injection site and slices were selected in which ∼10–50% of neurons were seen to be infected on the basis of GFP expression as assessed by eye before recordings. For paired whole-cell patch-clamp recordings, slices were transferred to the recording chamber with ACSF (127 NaCl, 25 NaHCO3, 1.25 Na2HPO4, 2.5 KCl, 2 CaCl2, 1 MgCl2, 25 glucose, saturated with 95% O2/5% CO2). Whole-cell patch clamp recordings were acquired simultaneously from neighboring Cre+ and Cre- pyramidal neurons in superficial CA1 and extracellular stimulation of local axons within specific lamina (SP or SR) of the hippocampus was delivered by current injection through a theta glass stimulating electrode that was placed in the center of the relevant layer (along the radial axis of CA1) and within 100-300 µm laterally of the patched pair. eIPSCs were pharmacologically isolated with CPP (10 μM) and NBQX (10 µM) in all experiments. Patch pipettes (open pipette resistance 2–4 MΩ) were filled with an internal solution containing (in mM) 147 CsCl, 5 Na2-phosphocreatine, 10 HEPES, 2 MgATP, 0.3 Na2GTP and 2 EGTA (pH=7.3, osmolarity=300 mOsm) and supplemented with QX-314 (5 mM). All recordings were performed at 31° C.

Electrophysiology data were acquired using ScanImage software ^77^ and a Multiclamp 700B amplifier. Data were sampled at 10 kHz and filtered at 6 kHz. Off-line data analysis was performed using NeuroMatic ^78^. Experiments were discarded if the holding current for pyramidal cells with CsCl-based internal solution was greater than −500 pA, if the series resistance was greater than 25 MΩ, or if the series resistance differed by more than 25% between the two cells. Individual traces were examined and if either recording contained spontaneous events that obscured the evoked IPSC then both the Cre+ and Cre- sweeps were excluded and average traces were created from technical replicates. Ratio-paired t-tests were performed comparing Cre+ to neighboring Cre-eIPSC amplitudes.

### Optetrode Fabrication

Optetrodes were fabricated following previously published designs with slight modifications ^79^. Briefly, the tetrodes used in the optetrodes were prepared by braiding four platinum-iridium wires (0.0007 mm diameter; California Fine Wire Company) together and applying heat to bind the wires together. Four of these tetrodes were then loaded into a 16-channel electronic interface board (EIB-18, Neuralynx) and pinned in place with gold pins to ensure stable connection with the EIB. An optic fiber (200 μm diameter; Doric Lenses; product code: MFP_200/240/900-0.22_#.#_SMA_ZF1.25(F)) was inserted through the middle of the four tetrodes such that the tetrodes evenly surrounded the optic fiber. The tetrodes were secured to the tip of the optic fiber using a small amount of glue before being plated with a platinum-iridium solution to achieve impedances between 100 and 200 MΩ.

### Optetrode Implantation

All surgeries were performed in accordance with (NIH) guidelines and following the approval of our protocol by UC San Diego’s IACUC. Optetrode implantation surgeries were performed on mice who had recovered well from the injection surgery (5-14 days between surgeries; 8 adult male mice). Animals were injected with a slow-release buprenorphine (0.02 mg/kg) subcutaneously pre-operatively which provided analgesia for 2-4 days post-op. Animals were anesthetized with isoflurane for the duration of the surgery (1.5%-2% isoflurane vaporized in oxygen) and body temperature was maintained at 37° C. Following three repetitions of cleaning the skin with betadine and 70% ethanol, the previous incision site was reopened and the skull exposed. Four stainless steel screws were anchored into the skull to provide stabilization and support for the implant. The same coordinates used for viral injection were used for the site of the craniotomy while a ground screw was inserted at the same AP coordinates in the left hemisphere. Following a craniotomy and durotomy, a stereotaxic frame was used to slowly lower the tetrodes into the brain. The tetrodes were lowered to a depth of ∼0.5 mm and the entire craniotomy was covered with gel (sodium alginate cured with calcium chloride) to protect any exposed brain and tetrode wires. The entire skull was then covered in dental cement to firmly secure the optetrode to the skull and anchor screws. Following surgery, the mice recovered in their home cage over a heating pad until awake and moving.

### Handling and Behavior

Once mice had recovered from the optetrode implant surgery (minimum of five days) we began habituation and food deprivation. Food deprivation was slowly introduced over 4-7 days until mice reached ∼90% of their full body weight. For the first three days of habituation, mice were brought to the experimental room and handled by the experimenter for 5-15 minutes. On days 4-6, 20-30 chocolate sprinkles (the reward used in the task) were randomly placed on the track and mice were allowed to forage for 15 minutes or until all the chocolate sprinkles were gone. Once mice ate 80% of the chocolate sprinkles within 15 minutes, we began task training.

The task consisted of a figure-8 maze that had the central arm blocked off so that mice could only run along the outer rectangular track. To begin with, mice were blocked into one arm of the track. Once data acquisition had begun, one of the blocks was removed (alternated each day) and the mouse was allowed to run in one direction around the track, receiving a chocolate sprinkle at the front center of the track, opposite of the starting point, for each trial. The second block was removed to allow running of full laps. At the end of each epoch of trials (5 trials for training and 10 for recordings), a block was placed just after the reward zone forcing the mouse to turn around and run the other direction for 10 trials. A recording session ended when the mouse had run 80 trials or for 30 min., whichever occurred first. If the mouse was unable to run 60 trials in 30 minutes, the session was excluded from analysis.

Before and after each track recording session, home cage recordings were obtained. Home cage recordings took place in the animal’s cage which was placed just to the side of the track and in view of the camera. If units were recorded that day, we also obtained an optostimulation recording in the home cage at the end of the session.

### Electrophysiology Recordings in Behavior

After the animals had recovered from the optetrode implant surgery (minimum of five days), the tetrodes were slowly advanced over the course of several days until the hippocampus was reached. CA1 was identified by the presence of strong theta oscillations in the LFP, ripples, and the presence of well-isolated clusters. Just before reaching CA1 and throughout the rest of the experiment, the tetrodes were lowered 14-28 μm per day to ensure that the recordings were stable and that new cells were recorded on a daily basis. Once all tetrodes left the CA1 pyramidal cell layer and had clearly entered stratum radiatum as reflected by inversion of ripples and lack of excitatory cell activity, no further recording sessions were performed.

To perform the recordings, the microdrive was connected to a digital Neuralynx recording system through a multichannel headstage preamplifier. The headstage and preamplifier were supported with elastics to assist the mouse in holding the weight. The LFP was band-pass filtered (0.1 to 8,000 Hz) and a threshold of 45-60 μV applied to isolate putative spikes. The LFP was continuously sampled at 32,000 Hz from one of the wires on each tetrode.

### Position Tracking

To track the animals’ position, we used a previously published, open access method ^80^. An Arduino (Mega 2560) was programmed to deliver a synchronizing pulse that consisted of 1 msec on, 1 msec off, followed by a series of pulses that counted up from 0 in binary. This pulse was fed into one of the CSC channels of the Neuralynx system and was also fed into the audio output of a camcorder (Sony HDR-CX380) that was used to obtain video of the animal’s position. The animal’s position was estimated to a high degree of certainty using LEDs that were mounted on the headstage preamplifier. If the position could not be obtained (primarily occurring when the preamplifier cord moved between one of the LEDs and the camera), a value of NaN was assigned. Using the pulse on the audio channel and the pulse on the CSC channel, a custom MATLAB workflow was generated allowing us to synchronize the animal’s XY position with the Neuralynx recordings. We validated the accuracy of this procedure as described in the initial publication^80^.

### Optotagging

At the end of each recording day, a 20–30-minute optotagging session was conducted. A laser (Opto Engine P/N:MBL-III-473-100mW) was used to deliver 473 nm wavelength light through a patch cable (SMA, 200 μm core, NA 0.63, Thor Labs) to the optic fiber in the optetrode assembly. Before the recording began, the light power was carefully set so that a small but discernible response could be observed occasionally in the LFP but no population spike was elicited (typically ∼0.3mW, **Figure S2A**). The population spike cluster was easily discernible by eye when it did occur as the amplitude was large and roughly equivalent for all the channels of a given tetrode (**Figure S2B**). For all optotagging experiments, light was delivered at 0.5 Hz; light-on for 10 msec.

For the high-power validation experiments, we waited until the end of the normal optotagging session then increased the laser power such that a population spike was noticeable on all four channels. During the high-power stimulation, we observed a clear response in the LFP and population spikes on all tetrodes on which units had been recorded that day (**Figures S2B and S2D**). In some cases, some of the clusters nearly disappeared suggesting that these were opto-tagged cells that contributed to the population spike. Following stimulation, all previously identified clusters reappeared. While we cannot directly prove that the unit identity was the same before and after light stimulation, we observed that the re-emerging clusters maintained consistent waveform features — including peak amplitude, energy, and peak-to-valley ratio — and remained in the same region of cluster space as before stimulation. These observations support the interpretation that the same neurons were recovered after high-power stimulation.

### Spike Sorting and Cluster Quality

The spike sorting software MClust (MATLAB 2009b, Redish Lab; https://redishlab.umn.edu/mclust) was used for spike sorting. Cluster cutting was performed manually using two-dimensional projections of the parameter space. For our cluster cutting parameters, we used amplitude, peak-valley ratio, and waveform energy. In most cases, the cluster cutting boundaries were originally established in the track recordings then applied to the home cage and optotagging recordings. To be included in analysis, all clusters had to appear qualitatively well-separated. Cluster quality was quantitatively assessed using the L-Ratio and Mahalanobis distance for units recorded on tetrodes with all four working channels (**Figure S2G and S2H**).

### Unit Classification

During cluster cutting, we immediately excluded clusters if they had a mean firing rate above 5 Hz as this would suggest that either this was an interneuron or it was an overlapping cluster of two cells. Although others have used other metrics (such as the shape of the waveform or the burstiness of the cell) to isolate interneurons we did not do that in this study for two reasons. First, the interneurons that are known to be involved in the NPAS4 phenotype are CCK basket cells^22^ which are regular-spiking cells that do not have a narrow waveform. Second, although CCK basket cells are not bursty and can be separated from pyramidal neurons using a burst index, we observed differences in bursting as part of the NPAS4 phenotype and could not use this as an exclusion metric. For these reasons, it is possible that our WT population may contain a small subset of interneurons^47^. Importantly, if this were the case, given what we know about these interneurons, it would obscure the differences between WT and KO neurons that we report here. We expect that our KO population is entirely excitatory since Cre is expressed under the CamKII promoter.

Next, the remaining population of cells was sorted into putative WT cells, KO cells, or excluded cells based on their optotagged response. For each cell, we separated the spikes that occurred during the optostimulation session into trials (duration of 2 s per trial). We aligned spikes according to when the light pulse was delivered and calculated the peristimulus time histogram (PSTH) using bin sizes of 1 ms. Using the PSTHs, the opto-response was defined as the maximum response when the light was off subtracted from the maximum response when the light was on. If a cell appeared ambiguous (i.e. was low firing or had an opto-response of 1) we excluded it from all analyses in this paper. Cells that had an opto-response greater than 1 (i.e. in which the maximum response in the PSTH during light-on was greater than the maximum response in the PSTH during light-off) were considered KO cells while cells with an opto-response less than 1 were considered WT cells.

### Perfusion and Tissue Processing in Behavioral animals

At the end of the experiment, all mice were anesthetized with a mixture of ketamine and xylazine (100 mg/kg ketamine, 10 mg/kg xylazine) and were perfused with ∼40mL of saline followed by ∼40mL of 4% paraformaldehyde (PFA). The tetrodes were carefully raised out of the skull and the brains were extracted and drop-fixed for an additional 24-48 hours in 4% PFA. The brains were rinsed in three 10-min washes with 1X PBS and left in a 30% sucrose solution for 24 to 48 hours or until the brains had sunk. A microtome was used to section the brains into 50 μm coronal sections which were then mounted onto slides, stained with DAPI, and coverslipped.

### Histology and Identification of Tetrode Locations in Behavioral Animals

All coronal sections spanning the dorsal hippocampus were imaged using a Keyence microscopy system. Images obtained at 2X magnification were used to confirm that the infection extended throughout dorsal hippocampus. Stitched 10X images were used to confirm the location of the tetrode tracts within the CA1 pyramidal cell layer. Finally, 60X images were obtained anterior to, posterior to, medial to, and lateral to the site of the tetrode implants to quantify the percentage of knockout cells (% GFP+; **Figures S1, S2I, and S2J**). All animals used in this study had infection that was primarily localized to CA1 (some expression in CA2 and cortex) and in which the implant site fully overlapped with the infection. Tetrode locations were primarily located medially in CA1 along the proximodistal axis with a slight skew towards distal CA1.

### Calculation of ISI Histograms and Burst Index

The interspike intervals (ISIs) were calculated by finding the difference in timing between each spike and the one after it. The ISIs were then binned into 10 ms bins and the burst index was calculated by taking the number of spikes that occurred at ISIs less than 10 ms and normalizing by the total number of spikes for that cell.

### Quantifying Firing Rates and Spatial Tuning

For each session, the velocity was calculated by averaging over 1 s using 0.1 s sliding windows. Only spikes during periods of running (velocity ≥ 2 cm/sec) were used for analysis. For each cell, we divided spikes into those that occurred when the animal was running clockwise and those that occurred when the animal was running counterclockwise, analyzing each set of spikes separately for all analyses. The mean and max. firing rates were obtained from the spatial maps and used to define the cut-off for low-firing cells. Cells with a mean firing rate < 0.1 Hz and a max firing rate < 1 Hz were excluded from analysis. Note that in a control analyses we excluded all cells with a mean firing rate < 0.5 Hz and a max firing rate < 5 Hz and the spatial tuning results still held. The track was linearized for each trial such that the reward zone was always at 0 cm and the center of the left and right arm were always 66 cm and 198 cm respectively.The animal’s position was binned into 4 cm bins, and the spike rate within each bin was used to construct raw linearized rate maps. These maps were then smoothed using a five-point symmetric weighted filter with weights [0.02, 0.10, 0.16, 0.10, 0.02], effectively approximating a Gaussian kernel. Bins in which the position could not be computed or the velocity was below the threshold were set to ‘NaN’ and appear blank or gray in the rate maps.

For the trial-averaged spatial tuning in Figure 2, we averaged across trials and used the trial-averaged maps to calculate the spatial tuning metrics (place field number, place field size, spatial information, and sparsity). To calculate the spatial tuning on individual trials, we first analyzed each trial individually and then used the average of all trial-wise values for each cell to statistically compare WT and KO populations. For analyses that depended on the identification of place fields (number of fields, size of fields, and all in-field/out-of-field analysis throughout the paper) a field was defined as the set of contiguous bins with firing rates above 10% of the max and in which one bin was greater than 50% of the max. For the average place field analysis, three neurons with very large place fields spanning the majority of the track were excluded. These units were suspected to be interneurons based on their atypically broad spatial tuning. Importantly, the statistical outcomes of the analysis were unchanged when these neurons were included. The signal-to-noise was the average in-field firing rate divided by the average out-of-field firing rate. The spatial information and sparsity were calculated as previously described^34^ using the equations 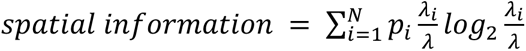 and 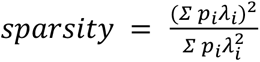 where 𝑖 = 1, …, 𝑁 represents the spatial bins, 𝑝 is the occupancy probability of bin 𝑖, 𝜆*_i_* is the mean firing rate for bin 𝑖, and 𝜆 is the overall mean firing rate for the cell.

### Stability of Firing Patterns Across Epochs, Shuffle Control, and Calculation of Place Field Shifts and Difference Maps

For all correlation analyses we used the Pearson’s correlation coefficient (PCC) on the trial-averaged rate maps. To compare firing in the clockwise and counterclockwise directions, we averaged the trials for each direction and calculated the PCC between them. To calculate the stability in spatial firing patterns across epochs we first averaged the trials for each epoch (set of 10 trials). We then calculated the PCC between each epoch and the subsequent epoch. For the shuffle control, we randomly shifted the rate map for each trial in space, enforcing a criterion that the shift must be at least 10 cm. We then averaged the trials for each epoch and calculated the PCC as described for the non-shuffled data. We repeated the shuffle 100 times and used the average PCC values for each cell. When calculating the stability using only in-field or out-of-field bins we used the trial-averaged rate maps for the full session and labeled bins as in-field if they were within a place field or out-of-field if not. We then repeated the stability analysis separately on the in-field and out-of-field bins. For all correlation calculations, if any bin had a value of NaN, it was removed from both vectors in that correlation comparison. To quantify place field shifts, we measured the change in the location of the peak firing bin between consecutive epochs for each place field independently. A negative shift indicated movement of the field toward the entrance of the track, while a positive shift indicated movement toward the field exit. Neurons with a peak shift less than −1 bin were classified as shifting toward the field entrance, those with a shift greater than +1 bin were classified as shifting toward the field exit, and neurons with a shift between −1 and +1 bins were considered stable.

Firing rate difference maps between epochs were computed by subtracting the firing rate map of the earlier epoch from that of the later epoch for each neuron (e.g., E2-E1, E3-E2, E4-E3). Difference maps were aligned by the E1 peak and displayed in the same neuron order across all comparisons.

### Analysis of the Local Field Potential (LFP) Using FOOOF (Fitting Oscillations and One-over-f)

Hippocampal LFP recordings from the last two track recording sessions containing units were analyzed across all animals. The LFP data was downsampled from 32 kHz to 1 kHz and preprocessed using the neurodsp package (https://neurodsp-tools.github.io/neurodsp/). This package was used to extract the frequency band of interest (0.1-100 Hz). Bandpass filtering was performed using a Butterworth filter provided through neurodsp (4th order filter, [0.1 - 100Hz]).

The FOOOF (Fitting Oscillations & One-Over F) package (https://github.com/fooof-tools/fooof) was used to analyze theta oscillations in the hippocampal LFP. This tool enables the characterization of neural oscillations by decomposing the power spectrum into a combination of periodic and aperiodic components. The theta frequency of interest was selected as 5-12 Hz. The FOOOF algorithm fits a model consisting of a combination of Gaussians to capture the periodic components (theta oscillations) and a smooth aperiodic function to describe the background activity. The parameterization process involved fitting the model to the power spectrum of each LFP recording. To quantify the power within the theta frequency band for each LFP recording, the power between 5-12 Hz was extracted and analyzed.

### Theta Modulation and Phase Precession of Single Units

For each cell, the LFP corresponding to the tetrode on which the unit was recorded was filtered in the theta frequency range using a Butterworth filter with cut-off frequencies of 4 and 12 Hz. For each spike a unit emitted, the theta frequency was obtained. Using circular statistics, we obtained the mean vector length and mean phase for each cell where 180° was the trough of theta and 0° was the peak. This same approach was used on spikes that occurred in-field/out-of-field and on spikes belonging to bursts/singles. For analyses separating bursts and single spikes, bursts were defined as spikes with interspike intervals (ISIs) less than 10 ms, and singles as spikes with ISIs greater than 10 ms.

For phase precession analyses, only in-field spikes were considered. For each trial, we plotted the theta phase of each in-field spike against the normalized position within the place field^37^. To account for the circular nature of phase data, we circularly shifted spike phases in 5° increments and computed the correlation between theta phase and spatial position at each step, identifying the shift that produced the maximum (most negative) correlation^54^. Using these optimally shifted phases, we fit a linear regression to describe the phase precession slope for each trial. To ensure sufficient data quality and reliable fits, we included only neurons with at least five in-field spikes per trial, a spatial span of at least three theta cycles, and a significant linear fit (*p* < 0.05). The median slope across trials was used to represent the overall phase precession for each neuron.

To assess the relationship between place field size and phase precession strength, we selected the trial with a slope closest to the median phase precession slope for each neuron. Place field size was log-transformed, resulting in an approximately linear relationship with phase precession slope. A simple linear regression was then performed, and we report the regression slope, R², and *p*-value. Theta modulation strength was defined using the mean vector length (MVL) as described above.

A multiple linear regression model was constructed using genotype, log-transformed place field size, and MVL as predictors of phase precession slope. Shuffling controls were performed by randomizing either spike phase or position within trials. Bootstrapping was used to test the robustness of slope estimates by randomly sampling neurons with replacement while maintaining the original sample size per genotype.

### Statistical Analysis

Statistical analyses were performed as described in the figure legends. Non-parametric tests were used unless otherwise indicated. Analyses were performed in GraphPad Prism and MATLAB.

### Use of generative AI tools

During the preparation of this work the authors used ChatGPT (OpenAI) to help revise and edit portions of the manuscript text for clarity and conciseness. No content was generated that affected the scientific results or conclusions. After using this tool/service, the authors reviewed and edited the content as needed and take full responsibility for the content of the published article.

## Supplemental Figures

**Supplemental Figure 1.**
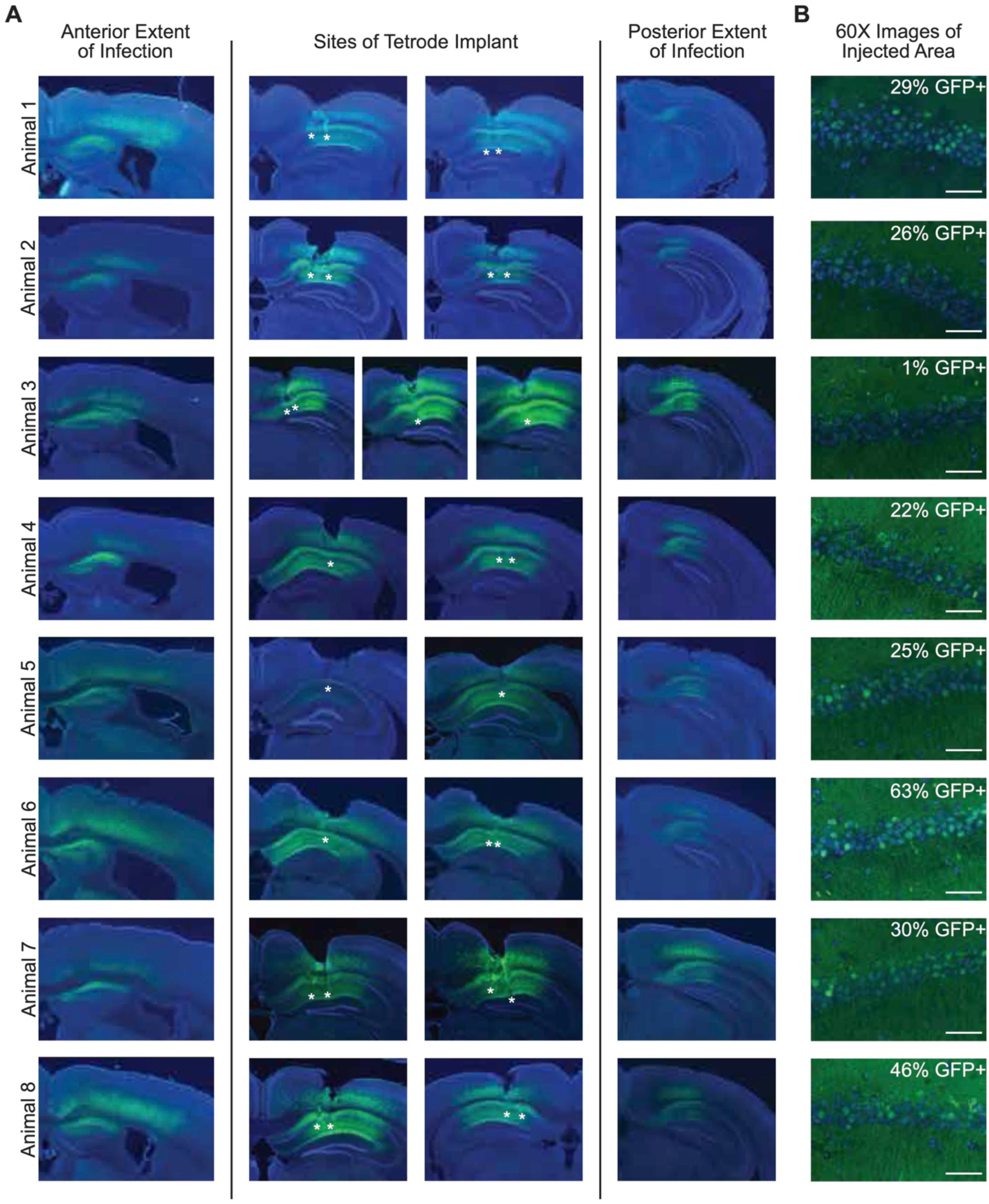
Representative histology for all sparse-infection animals. (A) Left: Stitched 10X images showing the anterior extent of the infection. Middle: Site of optetrode implant in CA1, asterisks represent the end of the tetrode tracts. Not all tetrode tracts were able to be identified. Right: Stitched 10X images showing the posterior extent of the infection. (B) Representative 60X images near the site of implant showing sparse infection. Scale bar = 50 µm.

**Supplemental Figure 2.**
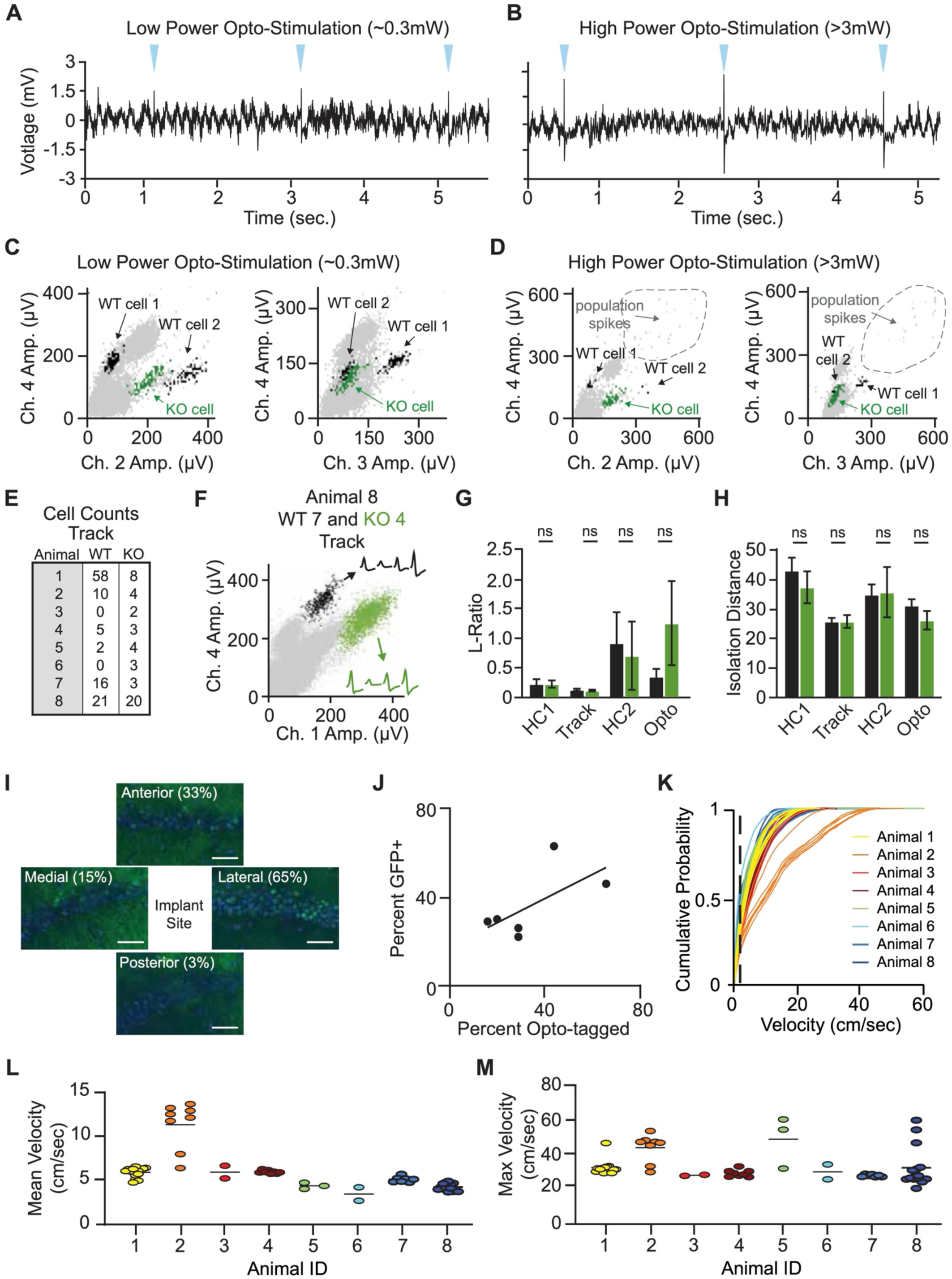
Optical identification and functional characterization of NPAS4 KO neurons in vivo. (A) Example of unfiltered LFP recording during light stimulation. Blue triangles = light delivery. (B) Example of unfiltered LFP recording during high-power light stimulation. (C) Example of cluster cutting for two WT neurons and a KO neuron during low-level light stimulation. Black dots: spikes from two WT neurons; green dots: spikes from one KO neuron. (D) Same neurons and cluster views as in (C) but for high-power stimulation. Dotted circle: putative population spikes. (E) WT and KO cell counts for each animal during track recordings. (F) Example of cluster cutting for a track recording showing a WT cell (black) and KO cell (green). Insets are average spike waveforms for each of the four tetrode channels for each neuron. (G) L-ratio for WT and KO cells for pre-track home cage (HC1), track, post-track home cage (HC2), and during optostimulation (HC1 WT N=70, KO N=34; track WT N=112, KO N=36; HC2 WT N=78, KO N=15; opto WT N=108, KO N=33; Kolmogorov–Smirnov test). (H) Isolation distance for WT and KO cells for HC1, track, HC2, and opto (HC1 WT N=71, KO N=36; track WT N=111, KO N=37; HC2 WT N=78, KO N=16; opto WT N=108 KO N=34; Kolmogorov–Smirnov test). (I) Example of how the percent infection is obtained from histology. 60X images taken anterior, posterior, medial, and lateral of the implant site. The percent infection per animal shown in (J) is an average of all four sites. Scale bar = 50 µm. (J) The percent of infected cells identified in histology plotted against the percent of optotagged cells identified *in vivo*. Line of best fit is shown. (K) Distribution of velocity for each session the sparse KO animals ran. (L) Mean velocity for all sparse KO animals. (M) Max velocity for all sparse KO animals. ns = not significant.

**Supplemental Figure 3.**
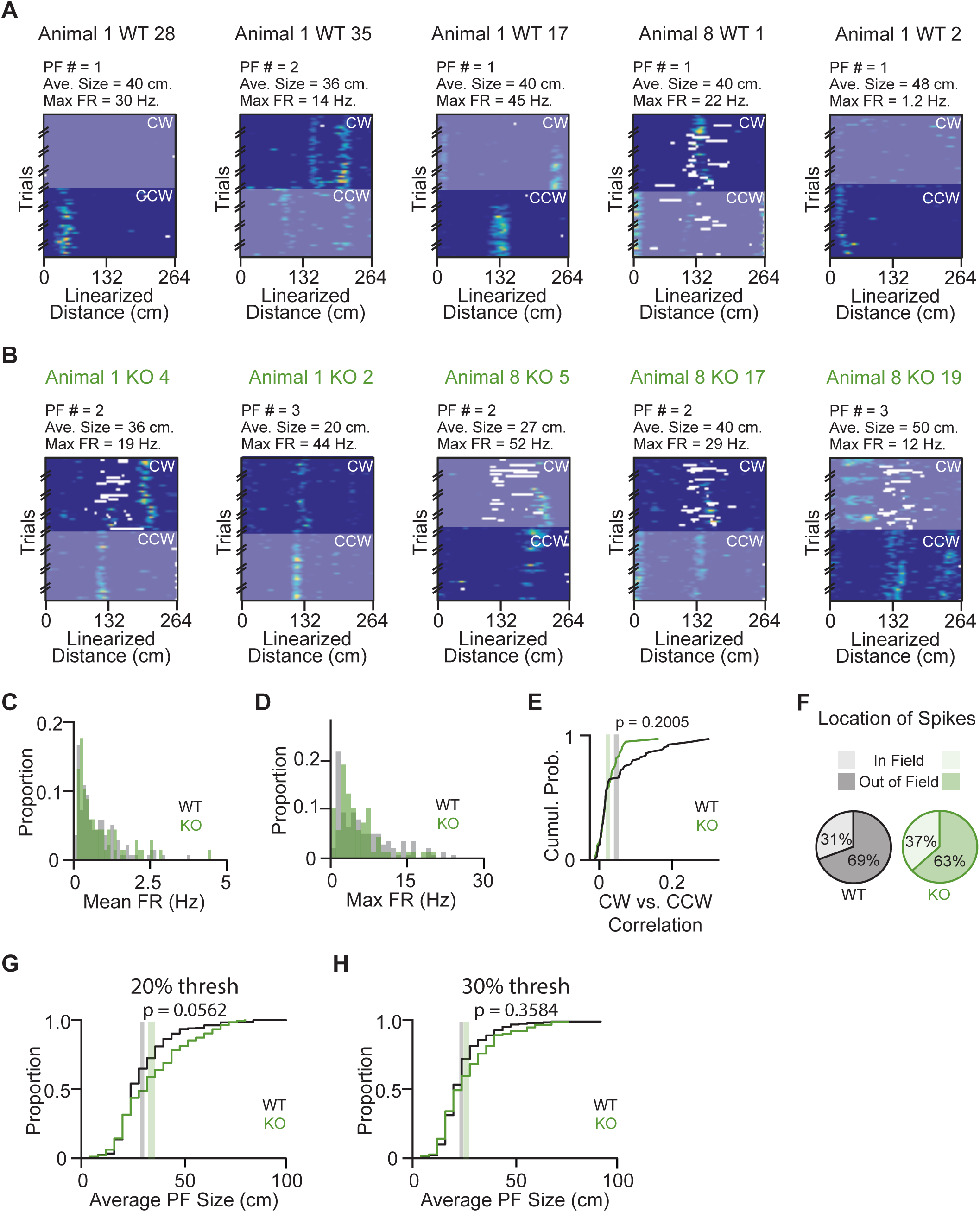
Additional information on spatial firing rate properties for NPAS4 WT and KO neurons. (A) Additional examples of rate maps from five WT cells. For each cell, both clockwise (CW) and counterclockwise (CCW) directions are shown; the direction not used for analysis is greyed out. The summary data shown above each rate map (Place Field Number [PF #], Average Size [Ave. Size], and Maximum Firing Rate [Max. FR]) corresponds to the analyzed direction only. (B) As in (A) but for KO cells. (C) Histogram of mean firing rate for all neurons (both low firing and high firing; WT: N = 224; KO: 94). (D) As in (C) but for the maximum firing rate. (E) Pearson’s Correlation Coefficient (PCC) between the trial-averaged CW rate map for each neuron and the trial-averaged CCW rate map (WT: N = 138; KO: N = 68; Kolmogorov–Smirnov test). (F) The percentage of all spikes for all neurons that occurred in-field and out-of-field for WT and KO neurons (WT: N = 140; KO: N = 68; p = 0.21; chi-square goodness-of-fit test). (G) Average place field size as calculated in the main figure with the exception that the minimum threshold used was 20% of max firing instead of 10% of max firing (WT: N = 138; KO: N = 68; Wilcoxon rank sum test). (H) As in (G) but with a minimum threshold of 30% (WT: N = 138; KO: N = 68; Wilcoxon rank sum test).

**Supplemental Figure 4.**
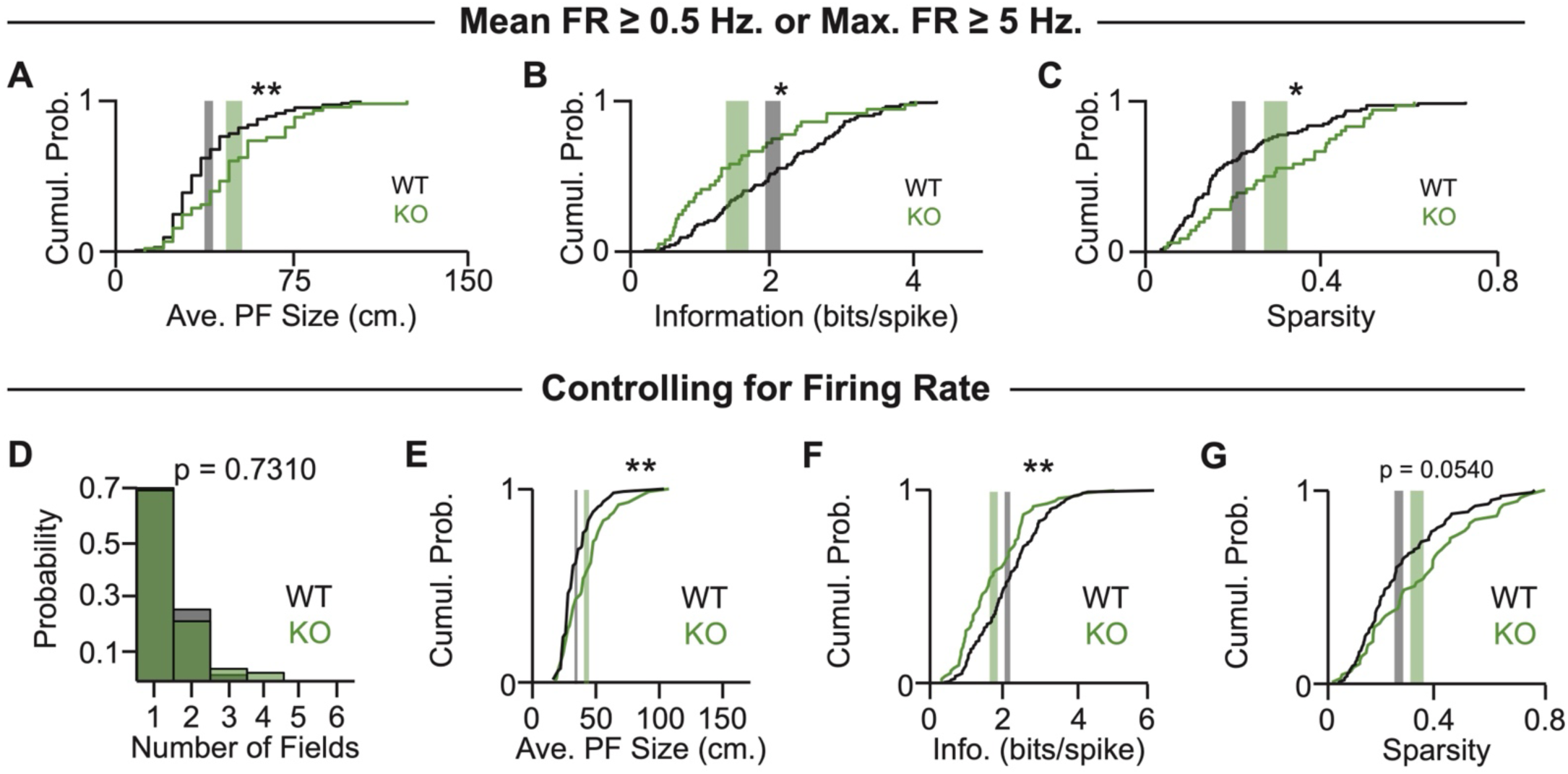
Spatial deficits persist across firing rate thresholds, matched firing rates, and independent replication. (A) Cumulative probability distribution of average place field size when only including neurons with mean firing rates > 0.5 Hz or max firing rates > 5 Hz. Gray shaded region: ± SEM for WT, centered at the mean; green shaded region: ± SEM for KO, centered at the mean (WT: N = 81; KO: N = 36; Kolmogorov–Smirnov test). (B) As in (A) but for spatial information. (C) As in (A) but for sparsity. (D) Histogram of the number of place fields per neuron when the number of spikes is the same across all neurons (WT: N = 138; KO: N = 68; Mann–Whitney test). (E) Cumulative probability distribution of average place field size when the number of spikes is the same across all neurons. Gray shaded region: ± SEM for WT, centered at the mean; green shaded region: ± SEM for KO, centered at the mean (WT: N = 81; KO: N = 36; Kolmogorov–Smirnov test). (F) As in (E) but for spatial information. (G) As in (E) but for sparsity. *p < 0.05; **p < 0.01.

**Supplemental Figure 5.**
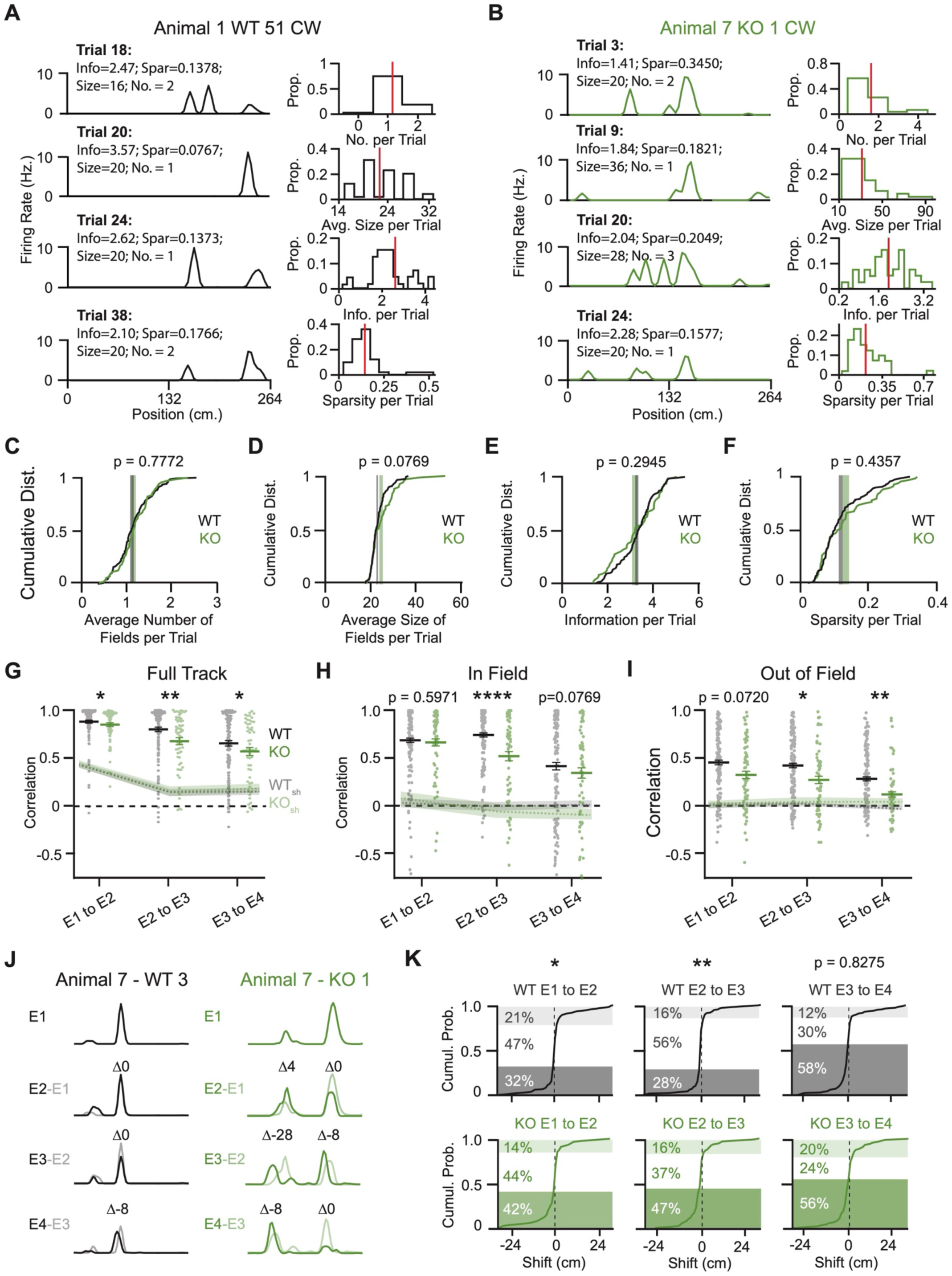
NPAS4 stability deficits are the result of spurious out-of-field firing and shifts in the place field towards field entrance. (A) Left: Four representative trials for a WT example cell. The number of fields, size, information, and sparsity are calculated independently for each trial. Right: For the same example cell, histograms for (top to bottom) the number of place fields, the average size of the place fields, the information, and the sparsity calculated for each trial independently and shown for all trials. Red line depicts the average. (B) As in (A) but for an example KO cell. (C) Median number of place fields per neuron. The number of fields was computed independently for each trial, then the median value was calculated for each neuron (WT N=140, KO N=68; Mann-Whitney test). (D) As in (C) but for the average place field size (WT N=140, KO N=68; Kolmogorov–Smirnov test). (E) As in (C) but for the spatial information (WT N=140, KO N=68; Kolmogorov–Smirnov test). (F) As in (C) but for the sparsity (WT N=140, KO N=68; Kolmogorov–Smirnov test). (G) Correlation between sequential sets of epochs (E1 to E2, E2 to E3, and E3 to E4) for all bins across the track. Gray dots: WT neurons; green dots: KO neurons; dotted gray: shuffled WT; dotted green: shuffled KO (WT N=140, KO N=68; Kolmogorov–Smirnov test). (H) As in (G) but for only the in-field bins for each neuron (WT N=140, KO N=68; Kolmogorov–Smirnov test). (I) As in (G) but for only the out-of-field bins for each neuron (WT N=140, KO N=68; Kolmogorov–Smirnov test). (J) Example trial-averaged rate maps for each epoch from one WT (left) and one KO (right) neuron to show shift calculation. Shift (denoted as Δ) is negative when the field shifts towards field entrance and positive when it shifts towards field exit. (K) Shift values for WT (top) and KO (bottom) across sequential sets of epochs (E2-E1, E3-E2, E4-E3). Dark shaded regions: fields with shift less than -1; unshaded regions: fields with shift between -1 and 1; light shaded regions: fields with shift greater than 1. Significance indicates comparisons between WT and KO (WT: N = 176 fields from 138 neurons; KO: N = 91 fields from 68 neurons). *p < 0.05; **p < 0.01; ****p < 0.0001.

**Supplemental Figure 6.**
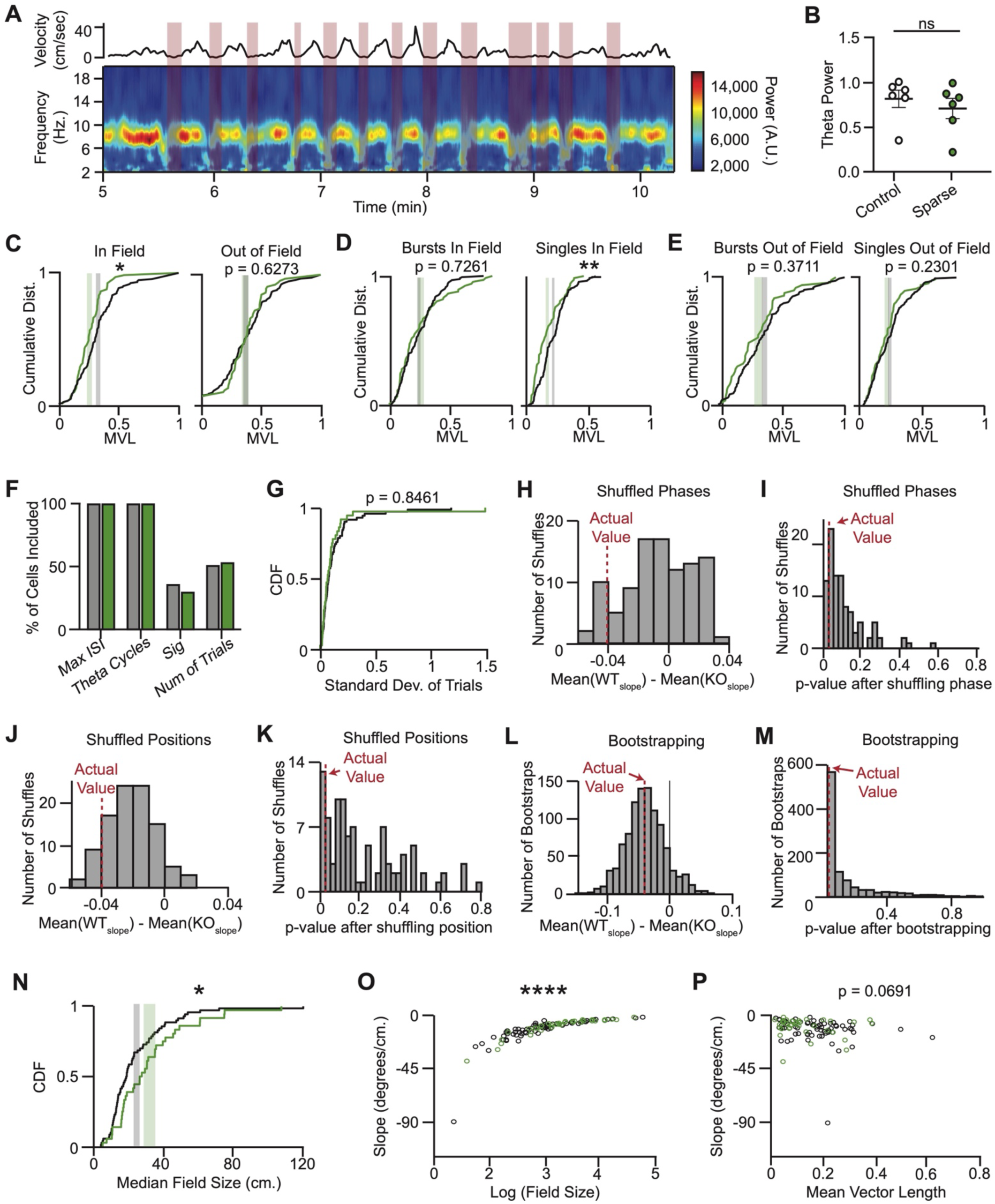
Impaired theta modulation in NPAS4 KO neurons is carried by single spikes but not bursts and the accompanying phase precession phenotype is related to differences in the size of the place fields. (A) Velocity (top) and spectrogram (bottom) for a representative session. Shaded red bars are periods of time when the velocity is below 2 cm/sec. (B) Theta power after accounting for the aperiodic offset (data are mean ± SEM; control N=6 animals, sparse N=8; Kolmogorov–Smirnov test). (C) Cumulative probability distribution of mean vector lengths for the in-field spikes (left) or out-of-field (right) spikes. Gray shaded region: ± SEM for WT, centered at the mean; green shaded region: ± SEM for KO, centered at the mean (WT: N = 138; KO: N = 68; Kolmogorov–Smirnov test). (D) As in (F) but for spikes in a burst that occurred in-field (left) or singles (spikes not in a burst) that occurred in-field (WT: N = 138; KO: N = 68; Kolmogorov–Smirnov test). (E) As in (F) but for spikes in a burst that occurred out-of-field (left) or singles (spikes not in a burst) that occurred out-of-field (WT: N = 138; KO: N = 68; Kolmogorov–Smirnov test). (F) Percentage of neurons retained after sequential thresholding steps applied during phase precession slope estimation. Criteria included: 1. Max ISI - spikes within each trial must have interspike intervals ≤ 1 s; 2. Theta Cycles - trials must span ≥ 3 theta cycles; 3. Sig - trial-level circular-linear regression must yield a p-value < 0.05; 4. Number of Trials - neurons must have ≥ 3 trials meeting the above criteria to be included in downstream analyses (WT: N = 70; KO: N = 36). (G) Cumulative probability distribution of the standard deviation of phase precession slopes across trials for WT and KO neurons. Gray shaded region: ± SEM for WT, centered at the mean; green shaded region: ± SEM for KO, centered at the mean (WT: N = 70; KO: N = 36; Kolmogorov–Smirnov test). (H-K) Shuffle analyses to assess whether the group difference in phase precession slopes reflects structured relationships between spike phase and position. (H) Histogram of mean slope differences (WT minus KO) across 100 iterations of theta phase shuffling, where theta phases were randomly permuted within each trial to disrupt spike–theta alignment. Red dotted line: the value derived from the actual data (WT: N = 70; KO: N = 36). (I) Histogram of p-values from Kolmogorov-Smirnov (KS) tests comparing WT and KO slopes in each phase-shuffled iteration (shuffling as in [H]). Red dotted line: the value derived from the actual data (WT: N = 70; KO: N = 36). (J) As in (H), but for position shuffling, in which spike positions were randomly permuted within trials (WT: N = 70; KO: N = 36). (K) As in (H), but for position shuffling as in [J] (WT: N = 70; KO: N = 36). (L-M) Bootstrapping analysis to estimate the reliability of the observed group difference. (L) Histogram of mean slope differences (WT minus KO) across 1,000 bootstrap iterations, resampling neurons with replacement while maintaining the original sample size per genotype (WT: N = 70; KO: N = 36). (M) Histogram of p-values from Kolmogorov-Smirnov (KS) tests comparing WT and KO slopes in each bootstrap iteration (WT: N = 70; KO: N = 36). (N) Cumulative probability distribution of the median field size for neurons included in the phase precession analysis. Gray shaded region: ± SEM for WT, centered at the mean; green shaded region: ± SEM for KO, centered at the mean (WT: N = 70; KO: N = 36; Kolmogorov–Smirnov test). (O) Relationship between phase precession slope and log-transformed field size across neurons (WT: gray; KO: green; WT: N = 70; KO: N = 36). (P) Relationship between phase precession slope and theta modulation strength (MVL; WT: gray; KO: green; WT: N = 70; KO: N = 36). *p < 0.05; **p < 0.01, ****p < 0.0001.

**Table S1.** Linear regression models examining predictors of phase precession slope. Models were fit using MATLAB’s fitlm function. Each model included different combinations of predictors: genotype (WT vs. KO), field size (log-transformed), and theta modulation (log-transform of the mean vector length). The purpose of these models was to determine which variables best account for variability in phase precession slope and to assess the unique contribution of each predictor. Adjusted R² values were used to compare model fit while penalizing for model

| Model | Predictors | Adjusted R<sup>2</sup> | Coefficients | p-Values |
| --- | --- | --- | --- | --- |
| 1 | Genotype only | -0.0050 | $\beta$ (intercept) = 1.8775<br>$\beta$ (genotype) = 0.0716 | p(intercept) = 3.6432e-43<br>p(genotype) = 0.6851 |
| 2 | Theta MVL only | 0.0323 | $\beta$ (intercept) = 2.2373<br>$\beta$ (theta MVL) = -1.7200 | p(intercept) = 1.8350e-31<br>p(theta MVL) = 0.0110 |
| 3 | Field size only | 0.2710 | $\beta$ (intercept) = -0.8943<br>$\beta$ (field size) = 0.9223 | p(intercept) = 0.0134<br>p(field size) = 2.4337e-13 |
| 4 | Genotype +<br>Field Size +<br>Theta MVL | 0.2784 | $\beta$ (intercept) = -0.6398<br>$\beta$ (genotype) = -0.2154<br>$\beta$ (field size) = 0.9191<br>$\beta$ (theta MVL) = -0.8995 | p(intercept) = 0.1149<br>p(genotype) = 0.1643<br>p(field size) = 1.3433e-12<br>p(theta MVL) = 0.1333 |
| 5 | Genotype *<br>Field Size +<br>Genotype *<br>Theta MVL | 0.2785 | $\beta$ (intercept) = -0.1919<br>$\beta$ (genotype, WT) = -1.3817<br>$\beta$ (field size) = 0.7841<br>$\beta$ (theta MVL) = -1.1387<br>$\beta$ (genotype, KO:field size) = 0.3894<br>$\beta$ (genotype, KO:theta MVL) = 0.2785 | p(intercept) = 0.7117<br>p(genotype) = 0.1036<br>p(field size) = 8.1668e-07<br>p(theta MVL) = 0.1081<br>p(genotype:field size) = 0.1613<br>p(genotype:theta MVL) = 0.7764 |

## Notes

### Competing Interest Statement

The authors have declared no competing interest.

